# Distinct neural geometries for target position and velocity in the primate superior colliculus

**DOI:** 10.64898/2026.06.22.733795

**Authors:** Feiran Yang, Clara Bourrelly, Mark A. G. Eldridge, Neeraj J. Gandhi

## Abstract

Moving targets pose a fundamental challenge for sensorimotor control: by the time a target’s position is transformed into neural signals, the target has already moved. Primates compensate for this delay and accurately intercept objects by accounting for target velocity. However, the neural mechanisms by which sensorimotor structures represent target velocity remain poorly understood. Here we recorded spikes and local field potentials from the superior colliculus (SC) of male macaques viewing or directing saccades to stationary and moving targets and used time-resolved dimensionality reduction and classification to characterize SC population dynamics. Target velocity emerged rapidly within the visual response and was robustly decodable despite weak and inconsistent single-neuron motion sensitivity. In low-dimensional state space, activity formed a V-shaped manifold in which speed was organized continuously along direction-specific arms. This geometry generalized across animals and signal modalities, could not be reproduced by displacement-based simulations, and remained largely dissociable from the axis encoding target location. Velocity information was strongest near the border between superficial and intermediate layers, identifying a laminar locus where visual motion signals may be transformed into commands for interception. These findings reveal separate, partially overlapping geometries for representation of target speed and position in the SC and overturn the longstanding view that it encodes only target location and movement goals.

**GRAPHICAL ABSTRACT:** 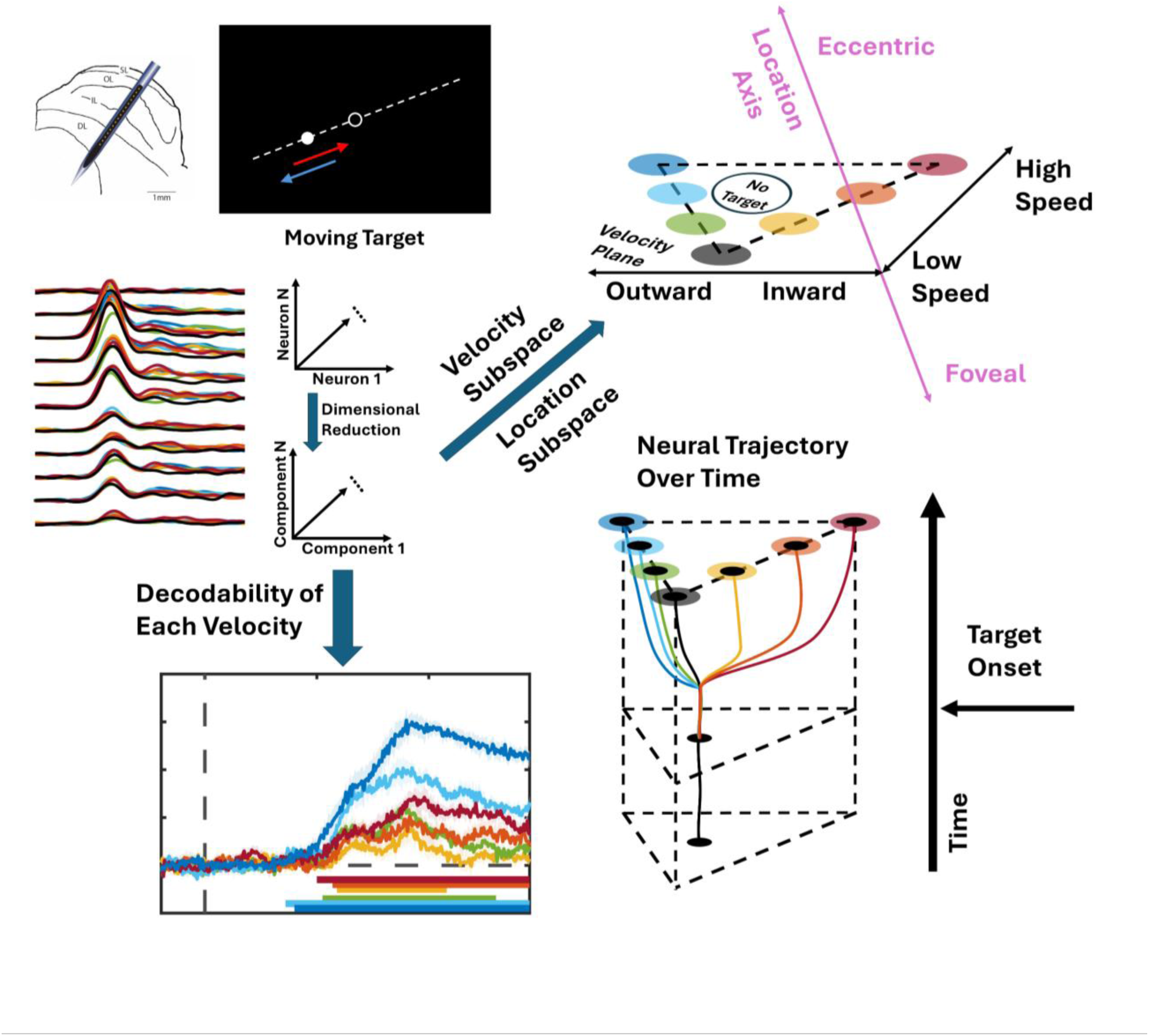

## INTRODUCTION

Intercepting a moving object requires the brain to solve a timing problem ^1–4^. Visual transduction, synaptic transmission, and sensorimotor integration impose delays ^5–9^, such that the neural estimate of target location available to downstream motor structures lags the true position. A location, or spatial, code alone is therefore insufficient for accurate interception. To overcome this delay, the brain must incorporate not only the lagged sensory location but also the velocity information to estimate the object’s current position. Indeed, many behavioral studies in humans, non-human primates and other species have demonstrated the ability to incorporate target velocity when generating interceptive movements ^10–21^. What remains unresolved is the neural basis of how motion signals are integrated in the circuitry that produces interceptive movements.

The superior colliculus (SC) is a compelling substrate for performing the computation. In primates, it is a subcortical hub for transforming visual information into motor commands for high-velocity eye movements known as saccades ^22–32^. It exhibits a laminar organization, with neuronal activity transitioning continuously from visually dominated responses in superficial layers to combined visual and motor activity in intermediate layers, and to motor dominated discharges in deeper layers. Though mostly studied in the context of stationary targets, the SC in sits at the converge of multiple motion-related inputs. Retinal signals can already carry predictive information about moving stimuli ^7,9,33,34^, and cortical motion pathways including the middle temporal (MT) area and frontal eye fields (FEFs) project to SC laminae that participate in sensorimotor transformation ^35–37^. This anatomy makes the SC well positioned not merely to inherit motion signals but to transform them into a format suitable for rapid orienting behavior.

Classical physiology in macaques, however, has left an apparent paradox. In several non-primate species, SC neurons are clearly tuned for motion direction or speed ^38–44^. In contrast, single neurons in monkey SC typically show weak, inconsistent, or stimulus-contingent motion selectivity, fostering the view that the primate SC is primarily a spatial orienting map rather than a substrate for motion computation ^25,45,46^. This conclusion, however, does not align with behavioral and physiological evidence that primates do not treat moving targets as static ones. SC neurons can signal motion in more complex visual contexts ^46,47^, and movement fields can reflect the executed interceptive saccade rather than the target’s onset position alone ^48^. These findings suggest that motion information is present in the primate SC but not in a form that is well captured by conventional single-neuron rate comparisons. We considered whether target velocity is embedded in population dynamics.

Across neural systems, latent population analyses have revealed computations that are weak or heterogeneous at the level of isolated neurons but become structured in multi-neuronal state space when analyzed across time and across channels ^49–52^. We asked if target velocity can be decoded from the SC population and, equally importantly, whether velocity is represented in a neural geometry that is distinct from the structure’s dominant spatial code. If so, the SC would need to be reconceived not only as a map of target location and saccade vector, but also as a site where dynamic motion variables are embedded within the population activity that links vision to action. Such a result would help explain how a structure classically associated with rapid orienting can support interception despite sensory delays.

We addressed these questions using simultaneous recordings of spikes and local field potentials (LFPs) from the SC of male macaques viewing or directing saccades to stationary and moving targets. We combined dimensionality reduction and classification analyses to determine whether target velocity is embedded in SC population dynamics on the timescale relevant for visual processing. We then tested whether any apparent velocity coding can be reduced to the trivial correlation between velocity and target displacement on a retinotopic map, using both behavioral tasks with opposite velocity-location relationships and simulations that model location coding in SC. Finally, we asked how the velocity representation relates to the axis coding target location and where along the SC laminar axis it is most strongly expressed.

We show that target velocity emerges rapidly within the SC visual response despite weak and inconsistent motion tuning at the level of individual neurons. In low-dimensional state space, SC activity forms a conserved V-shaped manifold in which velocity is organized continuously along direction-specific arms. This manifold remains largely distinct from the axis encoding target location and cannot be reproduced by position-only simulations. Velocity information is strongest near the superficial-intermediate transition, identifying a laminar locus where visual motion signals may be transformed into commands for predictive orienting. Together, these findings reveal a population-level velocity code in primate SC and suggest that a subcortical orienting structure participates in the computations that compensate for sensorimotor delay.

## RESULTS

Spikes and LFPs were recorded simultaneously from the SC of four monkeys performing Go and No-Go versions of a delayed saccade task to stationary and moving targets (Figure 1A-C; see Methods). Data were collected with laminar probes across 11 sessions in monkey DU, 11 sessions in Monkey SU, and 16 sessions in Monkey BU, yielding 264, 288, and 384 recording sites, respectively (Figure 1A). In monkey LO, we used a chronically-implanted, N-Form array with 16 shanks (4×4 layout, 8 recording sites on each shank; Figure 1B) designed to record neural data from 128 sites across the rostrocaudal, mediolateral, and dorsoventral dimensions within SC. This contrasts the laminar probe setup, in which the contacts only span the dorsoventral axis in a single penetration approximately orthogonal to the SC surface. However, spiking activity from the N-Form array was only available on the tip contacts of 12 of the 16 shanks. These neurons exhibited both visual and motor responses for targets and saccades in the upper left visual quadrant, indicating that the probe tips were positioned in the intermediate layers of the right SC. Data were collected across 9 sessions in monkey LO, yielding 144 recording sites. Spike sorting was performed on all recorded data (see Methods).

**Figure 1.**
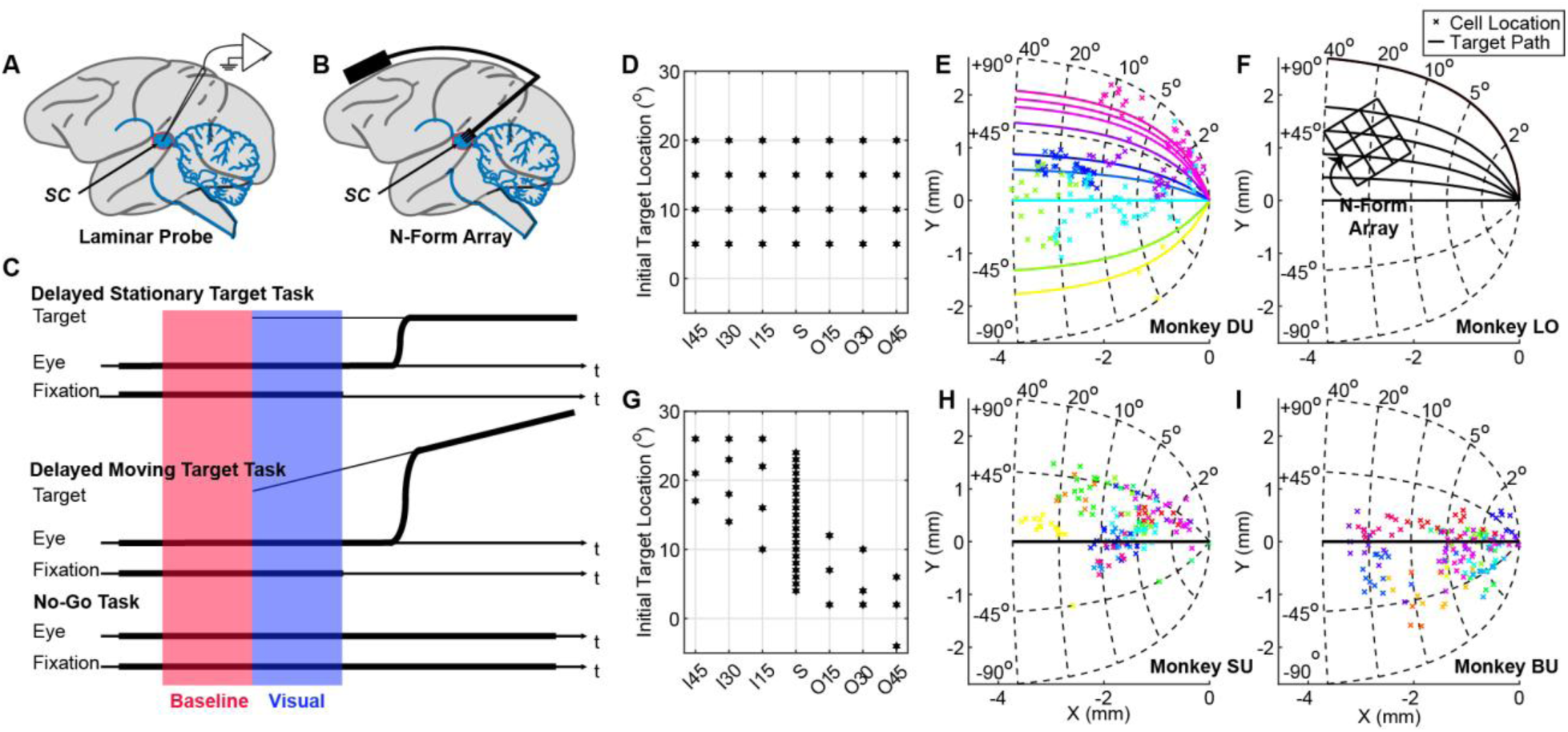
Recording setup and behavioral task paradigm. **(A)** Laminar probe recordings in three monkeys and **(B)** N-Form array recordings in one animal. **(C)** Schematic of the three task types: delayed stationary target task (top), in which a stationary target is presented after fixation; delayed moving target task (middle), identical to the delay stationary target task except that the target moves away from (shown) or toward (not shown) fixation; and no-go task (bottom), in which monkeys maintain fixation regardless of target status (not shown). **(D)** The same initial target locations were used for all speeds for monkeys DU and LO to ensure consistent spatial sampling across motion trajectories. **(E)** Monkey DU data were obtained from laminar probe recordings. Recording locations and target motion directions are projected on a topographic map of the SC. Each marker denotes the SC location associated calculated from the stimulation-evoked vector at each contact. The colored trajectory indicates the corresponding moving target’s path used in that recording session. Target directions were chosen such that moving stimuli passed through the estimated average receptive field center of the neurons recorded on a given track. **(F)** Monkey LO data were collected with an N-form array. The grid outlines the estimated array positions of the 16 shanks on the SC map; data were only available at the tips of the shanks. The black traces show topographic representation of target trajectories used across all sessions. **(G)** Initial target locations were not matched across speeds for monkeys SU and BU. The plot shows how initial target locations varied across stationary and moving target conditions. **(H & I)** The panels follow same convention as panel E. Data in both animals were collected with a laminar probe. Markers indicate recording sites and each color denotes a recording session. For these two animals, the targets moved only along the horizontal meridian. Thus, only one trajectory projection is shown.

Across all four monkeys, targets moved at three speeds in two directions (15°/s, 30°/s, or 45°/s; either inward toward fixation or outward away from fixation). Initial target placement was constrained by paradigm features like the delay period and task format (Go and No-Go) as well as the spatial operational range of the eye tracker (see Methods and Supplementary Table 1).

For monkeys DU and LO, target onset locations were matched across all target speeds and motion directions during the No-Go task. During the Go task, only the two most foveal target locations were used for outward moving targets, whereas only the two most peripheral target locations were used for inward moving targets (Figure 1D; see Methods and Supplementary Table 1). In contrast, for monkeys SU and BU, target onset location covaried with motion direction, with inward motion originating in the periphery and outward motion closer to the fovea (Figure 1G). In monkey DU, target trajectories passed close to the optimal location of the recording site (Figure 1E). In monkeys SU and BU, motion was constrained to the horizontal meridian regardless of track location (Figure 1H&I). In monkey LO, motion trajectories were presented along multiple direction axes in the upper left hemisphere to take advantage of the larger span of SC recorded with the N-Form array (Figure 1F, Supplementary figure 1).

### Single-Neuron Analysis Using Traditional Methods

To determine whether the neural responses of individual neurons were modulated by target motion, we first compared the activity levels elicited by moving targets with those evoked by stationary targets presented at the matching initial position. Figure 2 plots the spike density and LFP waveforms aligned on target onset for four initial locations (one per column) from an example session from monkey DU. Each trace represents one recording channel (arranged top to bottom for superficial to deep layers), with color indicating motion condition. Each waveform is an average across trials of matched conditions. The initial burst reflects the transient response to stimulus onset. Because all targets in each column appear at the same location, we did not expect differences in the initial visual burst between stationary and moving target conditions. The influence of motion emerged after this initial transient for both spikes (Figure 2A) and LFPs (Figure 2B). To quantify this effect across neurons and sessions, we selected two points for further analysis: the peak firing rate in the initial visual burst, whose timing varied slightly across monkeys, and the firing rate 180ms after target onset. For each speed and motion direction, we plotted and applied linear regression analysis to the trial-averaged firing rates for moving targets against those for stationary targets with the same initial location. For activity during the initial visual burst, data points clustered closely around the unity line in most cases (Supplementary figure 2, A-D, panel 1), indicating little to no difference between stationary and moving targets. In contrast, when comparing activity at 180ms after target onset (Supplementary figure 2, A-D, panel 2), we observed increased variance between stationary and moving targets.

**Figure 2.**
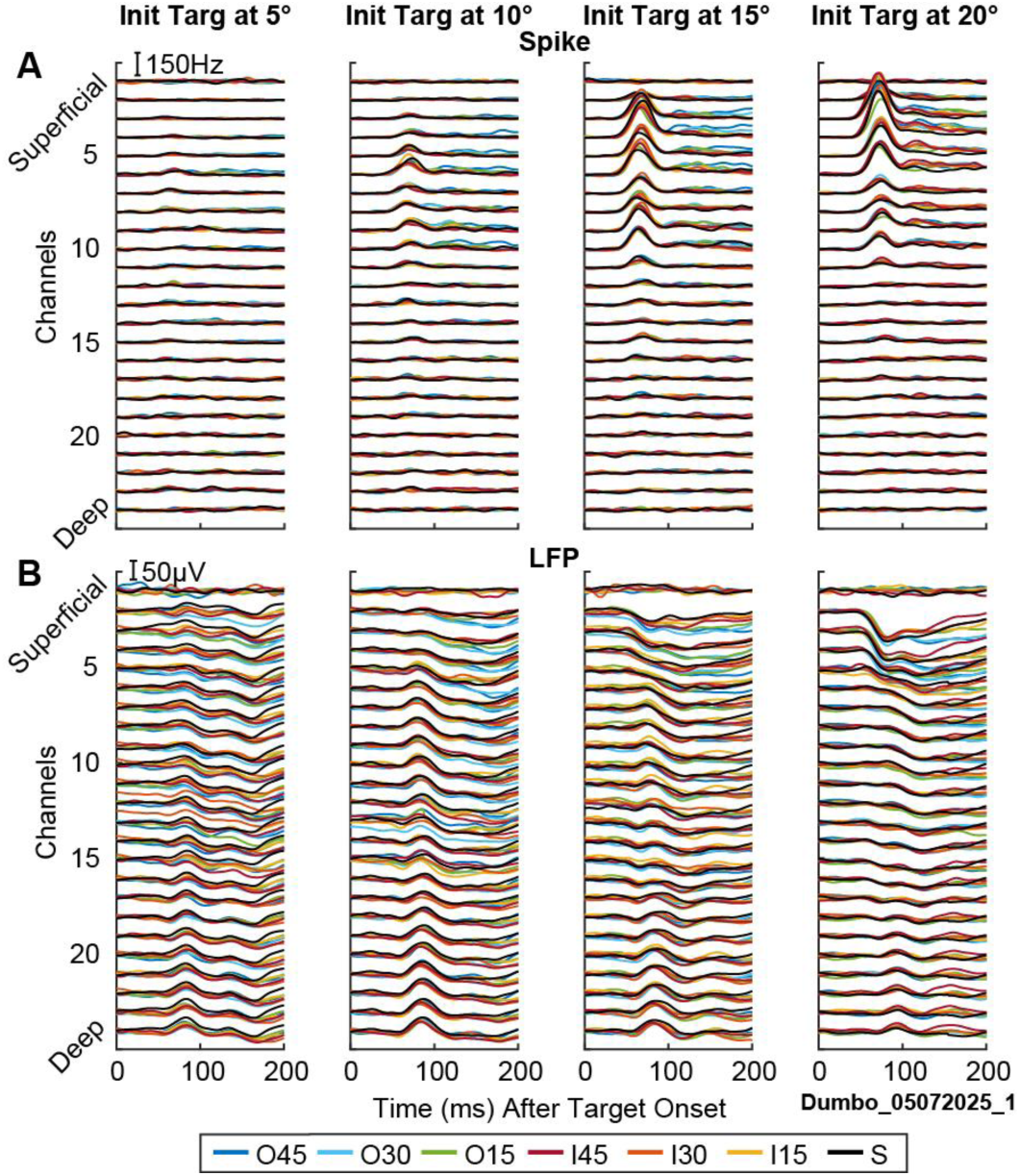
Single-neuron visual activity for moving and stationary targets. Example laminar recording session from monkey DU showing trial-averaged neural responses aligned to target onset. **(A)** Spike activity and **(B)** LFP activity are shown for four initial target locations (5°, 10°, 15°, and 20°; one column for each initial target location). Each row corresponds to one channel, ordered from superficial to deep along the probe. Within each channel, colored traces represent the seven target conditions: stationary targets and moving targets at three speeds in inward or outward directions.

To statistically quantify the separability between stationary and moving target trials, we computed receiver operating characteristic (ROC) curves comparing spike activity during the initial visual response (Supplementary figure 2A-D, panel 3) and at 180ms after target onset (Supplementary figure 2A-D, panel4). For each session, we performed the ROC analysis for each moving condition by pairing its individual trials with those from the corresponding stationary condition, matched for initial target location. ROC results were then aggregated across sessions to evaluate the reliability of motion-related differences in spike activity. Most AUC values clustered near 0.5 for spike activity at the initial visual response (Supplementary figure 2, A-D, panel 3), consistent with chance-level discrimination between the two groups. In some cases, significant differences were observed (asterisks denote p<0.05 for Benjamini-Hochberg corrected t-test). However, these effects were inconsistent across animals, conditions, or directions, and no systematic pattern emerged. For spike activity measured 180ms after target onset (Supplementary Figure 2, A-D, panel 4), AUC values began to deviate from 0.5, consistent with the increased variance observed between stationary and moving target responses. Nevertheless, this divergence was also not consistent across monkeys and yielded no systematic trend.

We also repeated the same analysis on the LFPs, both at its trough value (Supplementary figure 2A-D, panel 7) and at 180ms after target onset (Supplementary figure 2, A-D, panel 8). Overall, the distribution of LFP metrics was more variable than spikes. Though significant differences were observed at the trough and higher variability at 180ms after target onset, no consistent differences emerged across speeds, directions, or monkeys at either time window.

Taken together, these observations suggest that, based on traditional single-neuron analyses, both spikes and LFPs exhibit similar initial visual responses to stationary and moving targets, with divergence emerging only after target onset. Although differences were observed, they were inconsistent across sessions and monkeys, making it difficult to draw a systematic conclusion. Therefore, we turned to population-level analysis, an approach that minimizes redundancy and reduces noise across channels to capture subtle, temporally structured neural dynamics ^53,54^.

### Population-Level Analysis Using Dimensionality Reduction Methods

As illustrated in Figure 3, we applied principal components analysis (PCA) separately to the spike density or LFP waveforms recorded across contacts in each session. We retained the first 10 principal components (PCs) to account for at least 70% of the variance (Supplementary figure 3). We then segmented the low-dimensional neural activity into 20ms bins to capture population dynamics from 40ms before to 290ms (monkeys DU and LO) or 190ms (monkeys SU and BU; shorter epoch due to the shorter delay period) after target onset. Within each window, we tested whether these low dimensional features contained information about target motion using linear discrimination analysis (LDA) ^55^. We applied this method to train binary and multi-class decoders (Figure 3,①) to discriminate respectively between stationary and moving conditions or among the seven target velocity conditions (outward 45°/s, 30°/s and 15°/s; inward 45°/s, 30°/s and 15°/s; and stationary or 0°/s). Decoder performance and robustness were evaluated using Monte Carlo cross-validation with stratified sampling (80 % train, 20 % test, 100 iterations; see Methods). To obtain an estimate of chance-level performance for statistical comparison, we repeated the decoding procedures after randomly shuffling trial labels within each session.

**Figure 3.**
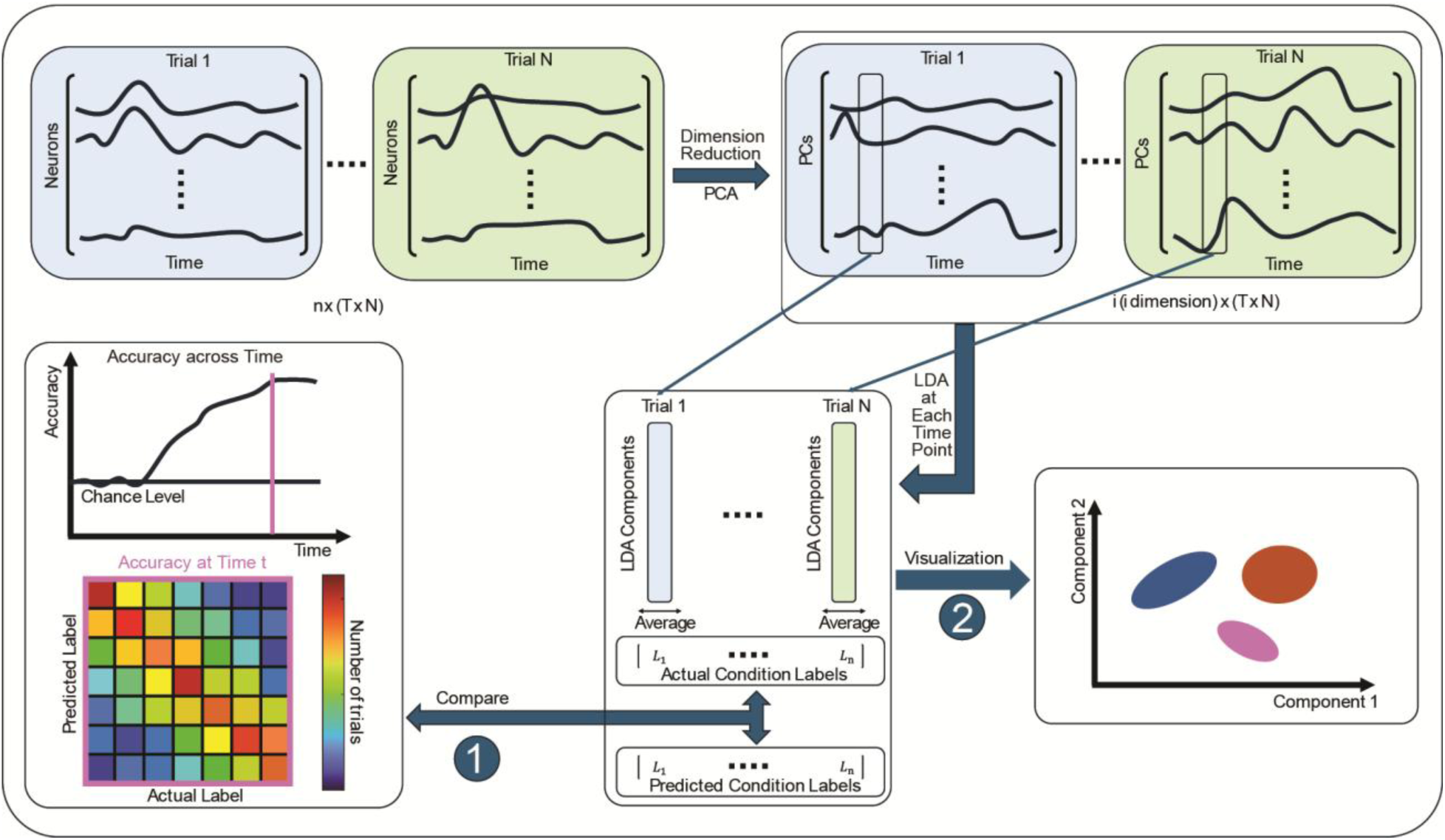
Illustration of dimensionality reduction method used for population-level decoding and visualization of target speed. PCA was applied separately to spike and LFP activity to reduce redundancy and noise. Trials were labeled according to either stationary or moving target for the binary decoder or seven speed categories (outward 45°/s, 30°/s, 15°/s; inward 45°/s, 30°/s, 15°/s; stationary 0°/s) for the multiclass decoder. Low-dimensional population activity was segmented into sliding time windows, and LDA decoders were trained to discriminate the category. Only the seven-speed condition is shown here **(①)**. The decoder operated on spike-derived PCs (“Spike”) or LFP-derived PCs (“LFP”). Decoder performance was evaluated at each time step using Monte Carlo cross-validation (80% train, 20% test, 100 iterations), and a confusion matrix was used to chart accuracy across speed categories. LDA components were also used to visualize each target condition in subspace representation **(②)**.

We first evaluated decoding accuracy when differentiating each velocity category from the stationary condition over time. Across all four animals, the binary decoder’s accuracy increased after target onset for each velocity category, followed by either a plateau or decline (Figure 4, panels 1 & 2 for each animal). Faster speeds (Pearson correlation; spike: p < 0.05; LFP: p < 0.01) and outward motion (paired t-test comparing inward and outward motion at the same speed within each monkey; p < 0.001 for both spikes and LFPs) consistently produced greater accuracy than slower speeds or inward motion (Supplementary figure 4A & B). The onset of significant decoding ranged from 20 to 120ms (horizontal lines below traces), depending on the monkey, velocity, and signal type.

**Figure 4.**
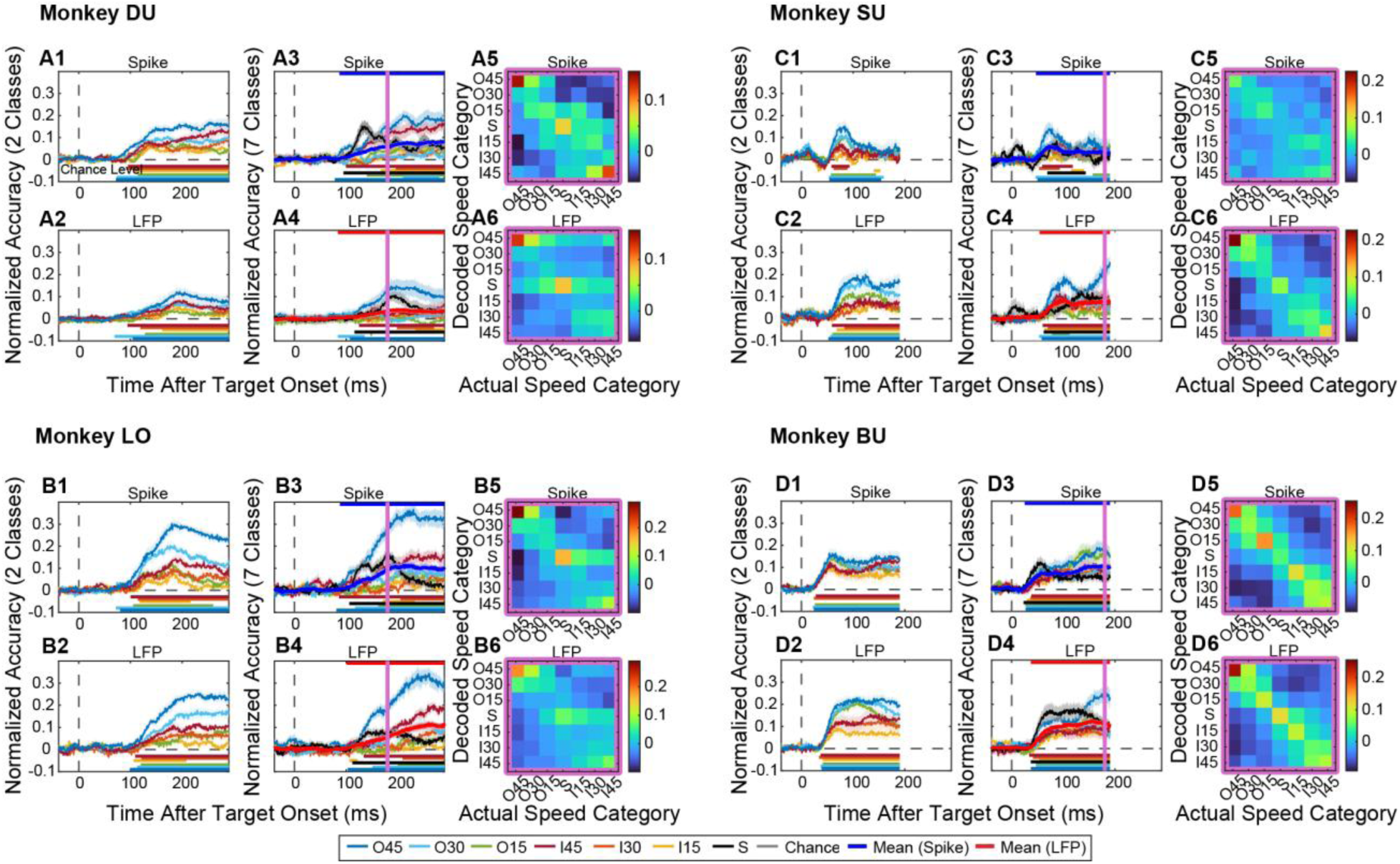
Time-resolved decoding of target speed averaged across sessions in each monkey. **(A-D)** Results for monkeys DU, LO, SU, and BU. **(panel 1-4)** Binary decoding accuracy **(panel 1 & 2)** distinguishing each moving target condition from the stationary condition for spike and LFP activity, and multiclass decoding accuracy **(panel 3 & 4)** across all seven velocity conditions. Normalized accuracy indicates calculated decoding accuracy minus shuffle accuracy. Target onset is indicated by the dashed vertical line. Shaded regions show mean ± SE across sessions. Dash line denotes chance level. Colored horizontal bars at the bottom of each panel mark the contiguous time clusters with decoding accuracy significantly above chance exhibiting the lowest p-value (cluster-based permutation test), while colored horizontal bars at the top in **panel 3 & 4** mark the significance for the averaged across categories decoding accuracy. Vertical purple lines indicate the time points used for confusion matrix analyses. **(panel 5 & 6)** Confusion matrices computed at the indicated time points for spike and LFP, averaged across sessions.

Multi-class decoding across seven velocity categories across the same timespan revealed a similar pattern (Figure 4, panels 3 & 4). Accuracy rose steadily after target onset and reached significance between 20 and 100ms (horizontal lines below traces; Spike: 87ms, 88ms, 47ms, and 25ms; LFP: 84ms, 101ms, 54ms, and 37ms; for Monkey DU, LO, SU and BU, respectively). This temporal profile paralleled the timing of visual bursts in the SC (Supplementary figure 4E & F), indicating that motion-related information emerges dynamically within the early visual response. The highest decoding accuracy was observed for outward 45°/s, inward 45°/s, and stationary conditions (Supplementary figure 4C & D). Mean accuracy exceeded chance for every signal type (mean ± standard error (SE); Monkey DU: Spike: 0.0580 ± 0.0045; LFP: 0.0359 ± 0.0072; Monkey LO: Spike: 0.0924 ± 0.0068; LFP: 0.0537 ± 0.0098; Monkey SU: Spike: 0.0280 ± 0.0117; LFP: 0.0820 ± 0.0133; Monkey BU: Spike: 0.1084 ± 0.0166; LFP: 0.1101 ± 0.0151; same cluster-based permutation test as above, p < 0.001). Control analysis revealed that eye drift, which was observed across animals and motion conditions, emerged substantially later than the onset of above chance decoding performance (Supplementary figure 5), implying that eye drift does not contribute to the observed velocity decodability.

Confusion matrices at 180 ± 10ms after target onset disclosed diagonal decoding patterns across all monkeys and decoders (Figure 4, panels 5 & 6), indicating above-chance correspondence between decoded and actual target velocities, particularly for the faster speed categories. Misclassifications were more likely to occur between neighboring velocity categories, suggesting limited but systematic velocity discriminability.

**Figure 5.**
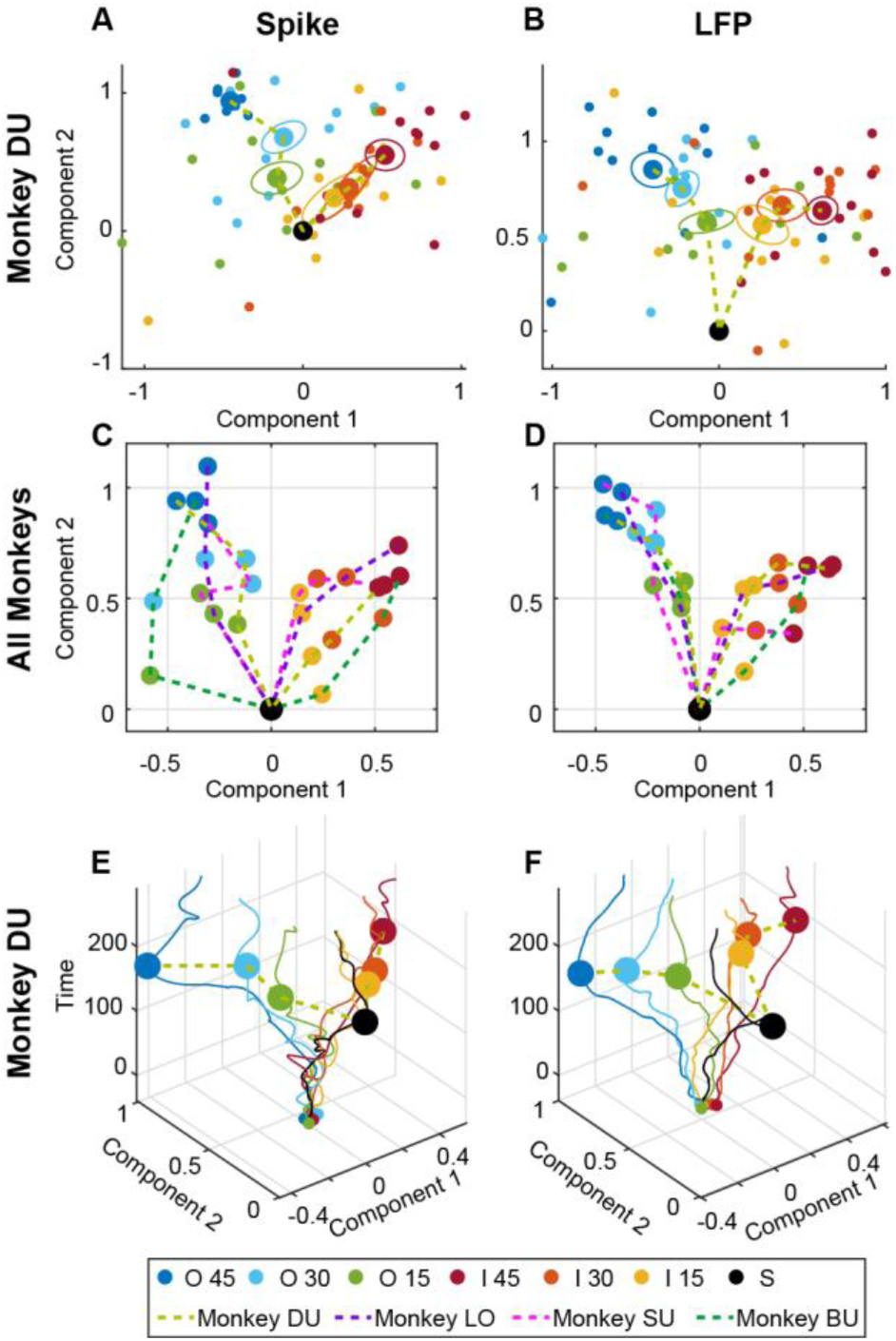
Low-dimensional subspace visualization of speed representations in SC population activity. **(A & B)** Spike and LFP activity from monkey DU are projected onto the first two LDA components at 180ms after target onset. Small dots indicate individual session centroids for each speed condition (outward 45°/s, 30°/s, 15°/s; inward 45°/s, 30°/s, 15°/s; stationary), color-coded by speed. Large dots represent the across-session mean centroid for each speed, and circles denote ± SE across sessions. The stationary data point (black point) is referenced to the origin, and the dashed line connects the big centroids in increasing speeds. **(C & D)** Across-monkey summary of mean centroids for spike and LFP activities at 180ms after target onset, averaged across sessions within each monkey. All four monkeys exhibit a similar V-shaped organization of speed representations in the low-dimensional subspace for spikes and LFPs. **(E & F)** Temporal evolution of the mean speed centroids is shown for spike and LFP activity for the same animal. Trajectories depict the population activity at each time window projected onto the two LDA components at 180ms (z-axis: time from target onset). Colors indicate speed condition. Large dots correspond to the mean location occupied at 180ms time point.

**Figure 6.**
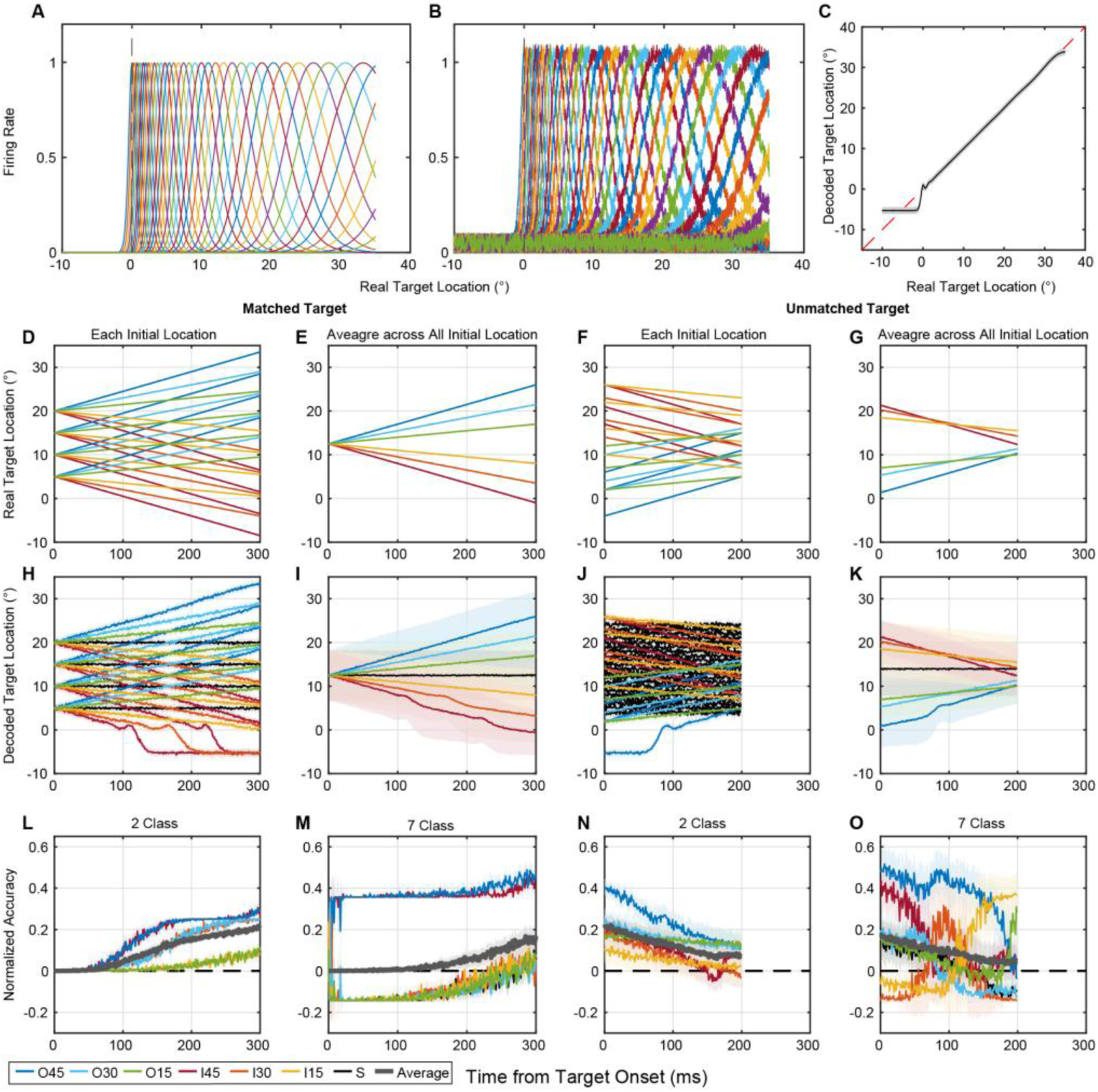
Simulated decoding based solely on target location. **(A)** Simulated neuronal receptive fields distributed uniformly and unilaterally across the SC map. **(B)** Simulated population responses after the addition of noise. **(C)** performance of the linear regression decoder used to estimate target location from population activity. The black trace shows decoded versus actual target location; shading indicates mean ± SD across simulated trials. **(D)** Temporal evolution of target location for the matched initial location condition. Each line represents one initial target location, and colors indicate target velocity. **(E)** Mean target trajectories averaged across initial locations for the matched condition. **(F & G)** Same analyses as in **(D & E)** for the unmatched initial location condition. **(H & I)** Decoded target location trajectories for the matched condition shown for individual target conditions **(H)** and after averaging across initial locations within each velocity condition **(I)**. Shading indicates mean ± SD across simulated trials. **(J & K)** Same analyses as in **(H & I)** for the unmatched condition. **(L-O)** Binary and multiclass decoding performance obtained from the simulated population activity for matched **(L & M)** and unmatched **(N & O)** target configurations. Colors denote velocity conditions, and gray traces indicate averages across conditions. Decoding accuracy is expressed as normalized accuracy (actual minus chance level).

We observed similar decoding performance for both binary and multiclass decoders across individual recording sessions before averaging across sessions. Likewise, confusion matrices exhibited consistent patterns. Supplementary figure 6 shows an example session, corresponding to the recording illustrated in Figure 2, including the LDA decoding results and associated confusion matrices.

Collectively, these analyses demonstrate that both spikes and LFPs in the SC carry consistent and extractable information about target motion velocity. Decoding accuracy increased rapidly after target onset, consistent with the timing of visual responses in the SC. Outward and higher-speed motions were decoded more accurately than inward or slower motions across monkeys.

### Representational Geometry of Target Velocity in SC Population Activity

Next, we sought to determine whether population responses to different target velocities form separable clusters and trajectories in state space. We therefore projected the first ten principal components into the first two LDA components to visualize population activity across different stimulus conditions (Figure 3,②). We also recentered, normalized and rotated each dataset’s LDA components to facilitate comparison across sessions and monkeys (see Methods section for details). Figure 5A & B plots the component values for the 180 ± 10ms window for all sessions in monkey DU. For both spikes and LFPs, the clusters formed a distinct V-shaped geometry in the LDA space (dashed traces): the stationary condition consistently anchored the vertex of the V, while increasing speeds extended along two diverging arms corresponding to inward and outward motion directions. Within each arm, clusters were organized progressively by speed, such that nearby speeds occupied adjacent positions and larger speed differences produced greater separations. This pattern was robust and consistent for both spikes and LFPs across all monkeys (Figure 5C & D).

We next examined how these cluster relationships evolved over time by projecting population activity at each time point onto the same subspace derived from the 180 ± 10ms window (Figure 5E & F). Before target onset, clusters overlapped closely, reflecting similar baseline activity. Following target onset, clusters gradually diverged, then formed the stable V-shaped structure described above. Similar temporal evolution patterns were also found in other monkeys (Supplementary figure 7).

To verify that the representational geometry was not confined to projecting the neural data on a timepoint long after motion onset, we also projected population activity onto velocity subspaces at different time windows (Supplementary figure 8). Across monkeys and signal types, population responses evolved from a weakly structured configuration before or shortly after target onset to a more organized low dimensional manifold at later time points, with the geometry becoming relatively stable after approximately 100ms. The V-shaped organization and separation between velocity conditions were largely preserved across subspaces defined at different time windows, indicating that the structure of the velocity related subspace is robust over time.

Together, these results demonstrate that the SC encodes target velocity within a continuous, low-dimensional manifold. The consistent V-shaped organization suggests that SC population activity represents velocity as a structured continuum rather than discrete categories. This geometric organization was stable across monkeys and signal modalities, revealing a continuous population-level sensorimotor representation.

### Accounting for the Inherent Correlation Between Velocity and Location

For smoothly moving stimuli, velocity and location are inherently correlated, with faster targets traversing larger distances than slower targets over the same time interval. Given the spatial dependency of SC receptive fields, differences in target location could potentially contribute to velocity decodability. We therefore evaluated whether the observed velocity decoding performance could instead be explained solely by target displacement.

The two experimental configurations used in this study generated opposite relationships between velocity and target location over time. In monkeys DU and LO, target onset locations were matched across velocity conditions (Figure 6D & E). As a result, location differences were on average absent at motion onset and gradually increased throughout the trial. In contrast, monkeys SU and BU were tested using unmatched onset locations (Figure 6F & G), producing the opposite pattern, where location differences were largest at motion onset and decreased over time. If velocity decoding was driven primarily by instantaneous target location, these two configurations should produce markedly different temporal profiles of decoding accuracy of velocity category.

To test this prediction, we simulated population activity that encodes target location but contains no explicit velocity information (Figure 6A-C; see Methods). Neurons were distributed across the

SC map unilaterally. Each unit’s receptive field is represented as a log-Gaussian in visual space (Figure 6A & B) or a Gaussian point-image in SC space. A linear regression model was used to estimate target location from population activity (Figure 6C & H-K). Since population activity is simulated only in one hemisphere of the SC, target locations in the opposite hemifield are weakly represented, leading to a discontinuity characterized by a step change from −10° to 0°. We then applied the same binary and multiclass LDA procedures to classify velocity categories from the decoded locations at each time bin (Figure 6L-O).

For the matched initial target location condition, the binary decoder gradually increased its ability to accurately differentiate between stationary and moving conditions (Figure 6L), qualitatively resembling the neural results (Figure 4A & B panel 1 & 2). However, the multiclass decoder yielded above chance performance only for the fastest inward and outward motion conditions (Figure 6M), in contrast to the neural data, in which all velocity categories were decodable (Figure 4A & B panel 3 & 4). For the unmatched initial target position data, the discrepancy was even more pronounced. For both binary and multiclass decoders, average classification accuracy decreased over time (Figure 6N & O), opposite to the temporal evolution observed in the neural data (Figure 4C & D). For the multiclass decoder operating on simulated data (Figure 6O), individual velocity categories exhibited inconsistent temporal profiles, with decoding accuracy increasing for some conditions and decreasing for others. In contrast, the neural data showed a stereotyped pattern in which decoding accuracy increased following target onset and then plateaued across all velocity categories (Figure 4C & D panel 3 & 4).

To match the spatial sampling of our recordings, we repeated the simulations using smaller neuronal populations distributed over restricted regions of the SC map (Supplementary figure 9). Under these conditions, decoding performance depended strongly on the sampled location and on whether target onset locations were matched or unmatched. In contrast, the neural decoding results were qualitatively similar across monkeys despite differences in target configurations and recording locations.

Together, these findings indicate that differences in target displacement alone cannot account for the observed velocity decodability. Instead, SC population activity contains information related to target velocity beyond that predicted from instantaneous target location.

### Separability of Target Motion and Position Representations

Having established that SC population activity encodes target velocity independently of target position and knowing that the neural activity is also sensitivity to target position, we next explored the separability of their representations. We previously identified a subspace that maximally separated velocity categories. Here, we identified a subspace that best represents target location and then quantified the relationship between these two subspaces.

To define the target location subspace, the same PCA reduced neural activity from a 70 ± 10ms window following target onset was projected onto the initial target location of each trial using linear regression. This time window corresponds to the peak visual response across monkeys and is commonly used for receptive field analyses. Because initial target location varied continuously, linear regression provided a natural mapping between neural activity and spatial position. The resulting regression coefficients were taken as the transformation matrix and defined the target location axis.

To evaluate how well this target-location axis captured spatial information throughout the motion trajectory, trials from all sessions were grouped according to their initial target location and velocity. Neural activity was averaged within each group and projected onto the target-location axis at each time point. We then compared the projected target-location coordinate with the actual target location 70ms before the projection time point to account for visual transduction delay. Results are shown for spikes (Figure 7A, E, I, & M) and LFP (Figure 7B, F, J, & N). For both matched and unmatched target configurations, we observed a clear and approximately linear relationship between the projected target-location coordinate and the actual target location over a range of approximately 5°-20°. This relationship was present across all monkeys, initial target locations, and velocity conditions. These results indicate that although the target-location axis was derived from neural activity shortly after target onset, it remained stable throughout the visual response and continued to accurately represent target location as the target moved. Deviations from linearity became more apparent for target locations below 5° or above 20°, where spatial coverage was limited. Similar distortions were observed in simulations using restricted neuronal populations sampled from the middle portion of the SC map (Supplementary figure 9). We therefore attribute these deviations primarily to incomplete sampling of the SC, particularly the absence of neurons representing more peripheral locations or locations represented in the opposite hemisphere. In the unmatched-target condition (Figure 7I, J, M, & N), the projected location estimates were noticeably noisier for stationary targets. We believe this reflects the design of the experiment rather than a limitation of the neural representation. Because the unmatched-target condition contained a larger range of initial target locations while maintaining a similar number of trials per velocity condition, substantially fewer trials were available for each individual initial-location group. The resulting reduction in trial count likely contributed to the increased variability observed in these conditions. Taken together, these results demonstrate that the target-location axis provides a robust representation of target position and closely tracks the true target location throughout the visual response period.

**Figure 7.**
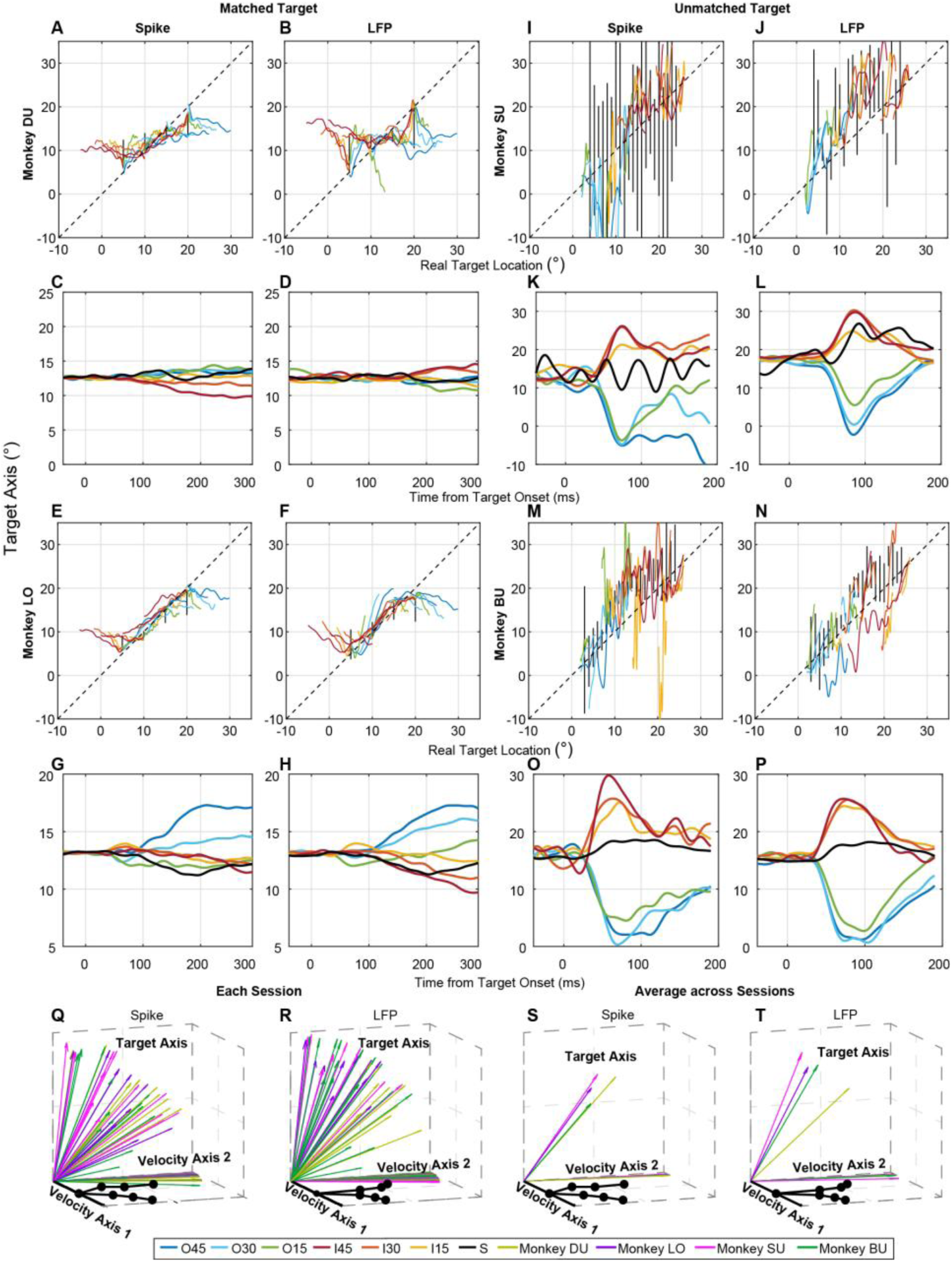
Low-dimensional representation of target location across matched and unmatched initial target conditions and its relationship to velocity encoding. (**A-H**) Matched-target analysis. Relationship between the projected target location based on spike (**A & E**) and LFP (**B & F**) and the actual target location 70ms before the projection time point (to account for visual transduction delay) for monkeys DU and LO. Colored traces represent individual velocity conditions. The multiple traces of the same color correspond to different initial target-location groups for the same velocity condition. The decoded target position is now plotted as a function of time for spike (**C & G**) and LFP (**D & H**) data, but data are averaged across initial target-location groups. (**I-P**) The same analysis is repeated for unmatched-target conditions used in monkeys SU and BU. (**Q & R**) Geometric relationship between the target-location axis and the first two velocity axes for individual recording sessions. Velocity axes were defined by the first two LDA components (Figure 5), whereas the target-location axis was defined by linear regression. To facilitate visualization of the seven velocity categories, the characteristic V-shaped manifold in the velocity plane was constructed from the session- and monkey-averaged velocity representations shown in **Figure 5C & D**. Each colored vector corresponds to a ngle recording session, with colors denoting monkeys. Spike activity is shown in **Q** and LFP activity in **R**. (**S & T**) verage relationship among the target-location axis and the first two velocity axes across recording sessions within ch monkey. Colored vectors denote monkey-specific averages. Same V-shaped manifold in the velocity plane as **Q &** Spike activity is shown in **S** and LFP activity in **T**.

We next projected neural activity across time onto the target location axis and examined its temporal evolution. Although activity was projected onto the same target location axis, this time trials were grouped by velocity condition, allowing us to assess the extent to which velocity related structure was present within the location representation. In monkeys with matched initial target locations (DU and LO; Figure 7C, D, G, & H), trajectories associated with outward motion shifted toward more eccentric positions after target onset, whereas inward motion trajectories shifted toward foveal position, consistent with the physical motion of the targets (Figure 6E) and our simulation (Figure 6I). However, substantial overlap remained between several velocity conditions. For example, stationary and outward motion trajectories overlapped extensively in monkey DU, whereas stationary and inward motion trajectories overlapped in monkey LO. This limited separation contrasted with the clear segregation of the same conditions within the velocity subspace (Figure 5E & F; Supplementary figure 7A & B).

A similar pattern was observed in monkeys with unmatched initial target locations (SU and BU; Figure 7K, L, O, & P). Trajectories initially reflected differences in initial target eccentricity and subsequently converged toward the center as target locations became more similar over time, consistent with the target motion itself (Figure 6G) and simulation (Figure 6K). Despite these pronounced location differences, separation among speed conditions within each motion direction remained weak along the location axis, again contrasting with the robust separation observed within the velocity subspace (Supplementary figure 7C-F).

Together, these results indicate that target velocity and target location occupy different subspaces within SC population activity. Although the two representations are not fully independent, the limited velocity related separation observed within the location subspace, compared with the strong segregation observed within the velocity subspace, supports the conclusion that SC population activity contains a distinct representation of target velocity.

Next, we examined the relationship between the velocity axis derived from the LDA subspace and the initial target location axis. The location axis was defined by linear regression coefficients, and the velocity axis was defined by the first two LDA components used for visualization. We computed the angles 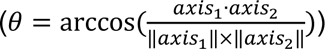 between each velocity axis and the location axis to assess their geometric relationship. Across monkeys, the two subspaces were not fully orthogonal but partially overlapping, with angles ranging from approximately 50° to 80° (Figure7Q & R, mean ± standard deviation (SD); Velocity Axis 1 VS Location Axis: Monkey DU Spike: 63.6 ± 7.7⁰; LFP: 58.6 ± 13⁰; Monkey LO Spike: 49.6 ± 16.8⁰; LFP: 60.2 ± 17.8⁰; Monkey SU Spike: 58.7 ± 13.9⁰; LFP: 70.1 ± 19.3⁰; Monkey BU Spike: 41.7 ± 19.2⁰; LFP: 63.8 ± 24.7⁰; Velocity Axis 2 VS Location Axis: Monkey DU Spike: 56.8 ± 14.8⁰; LFP: 55.6 ± 10.9⁰; Monkey LO Spike: 71.7 ± 12.7⁰; LFP: 74.6 ± 12⁰; Monkey SU Spike: 68 ± 17.6⁰; LFP: 71.4 ± 14.6⁰; Monkey BU Spike: 73.3 ± 11.3⁰; LFP: 68 ± 14.4⁰). Note that the vectors obtained from LDA are not constrained to be orthogonal. Therefore, the angle between the first and second velocity axes is only approximately 90⁰ (Figure7Q & R, mean ± SD; Velocity Axis 1 VS Velocity Axis 2: Monkey DU Spike: 82.7 ± 6.6⁰; LFP: 84 ± 5.4⁰; Monkey LO Spike: 84 ± 4.7⁰; LFP: 85.8 ± 3⁰; Monkey SU Spike: 86 ± 2.8⁰; LFP: 75.9 ± 6.2⁰; Monkey BU Spike: 83.6 ± 7.5⁰; LFP: 82.7 ± 5.6⁰). Figure 7S & T shows the relationship between the velocity axis and the target location axis averaged across sessions within each monkey. A similar organization was observed across all monkeys, with the velocity and location subspaces exhibiting partial overlap but remaining far from complete alignment.

These results indicate that although target location and target velocity are inherently correlated, a substantial degree of independence remains between the two features. This partial independence supports the interpretation that the velocity related subspace captures speed-related features rather than encoding simply target location information.

### Effect of Intracollicular Organization on Velocity Decodability

The SC contains a well-organized retinotopic map and a continuous laminar structure that transitions from visual dominant to motor dominant activity. Thus far, we have examined whether SC neurons encode target velocity in general. We next asked how multiple anatomical and functional features of SC organization, including neuron spatial location, motion direction, relative target position, and laminar depth, influence the decodability of target velocity. This approach allowed us to assess how structural and functional properties shape velocity representation. For neuron spatial location, motion direction, and relative target position, decodability was quantified within the window of 180 ± 10ms after target onset. For laminar depth, decodability was computed over the full interval from 0 to 180ms after target onset. Within the same layer, rostral SC neurons typically have smaller receptive fields and denser spatial representation, whereas caudal neurons have larger receptive fields and sparser populations. We therefore tested whether neurons located at different rostrocaudal positions differ in their ability to encode velocity. Horizontal cell location, defined along the x-axis in Figure 1E, F, H, & I, was used to represent relative rostrocaudal position. In laminar probe recordings, electrode penetrations were placed approximately perpendicular to the SC surface, such that neurons recorded within the same session preferred similar horizontal and vertical components. Each neuron’s location in SC was determined based on its preferred vector, and the location of each session was defined as the average position of all neurons recorded in that session.

Across the three monkeys recorded with laminar probes, only one animal showed a significant correlation between decoding accuracy and horizontal position (Supplementary figure 10C; Pearson correlation, p < 0.05 for both spikes and LFPs). No consistent relationship was observed in the other monkeys (Supplementary figure 10A & B), suggesting that rostrocaudal position alone does not systematically determine velocity decodability.

In the monkey with an N-Form array, similar (but not identical) neuronal population was sampled across sessions while the moving target followed different trajectories. This design allowed us to test whether SC neurons exhibit directional preference in velocity decoding. We found no significant correlation between decoding accuracy and motion direction (Supplementary figure 10D), indicating that target directional did not strongly influence decoding performance at the population level.

The relative spatial relationship between a neuron’s receptive field and the moving target trajectory is also likely to influence decoding accuracy. In most cases the direction of target motion did not align with the neuron’s preferred direction. We therefore examined whether angular misalignment between the motion trajectory and the receptive field center predicted decoding performance (Supplementary figure 10E). Across the three monkeys recorded with laminar probes, we observed a general negative relationship between angular misalignment and decoding accuracy, such that larger angular differences were associated with lower decoding accuracy (Supplementary figure 10F-H). However, this relationship reached statistical significance only in the LFPs of one monkey (Supplementary figure 10G; Pearson correlation, p < 0.05). To further evaluate this effect, neurons were grouped into near and far categories based on angular distance less than or greater than 25 degrees. Mean decoding accuracy was higher in the near group than in the far group (Supplementary figure 10I-J), with significant differences observed in several monkeys and signal types (t test; DU: LFP p < 0.05; BU: spike p < 0.005; SU: spike p < 0.05; SU: LFP p < 0.001). Together, these results suggest that spatial proximity between the motion trajectory and the receptive field enhances velocity encoding, although the effect varies across animals and signal types.

We next asked whether the initial proximity between the target onset location and the neuron’s receptive field center also influences decoding performance. To isolate this relationship from trajectory misalignment, we projected the averaged session location onto the motion trajectory axis to obtain a measure of relative distance between target onset and preferred location (Supplementary figure 10K). This analysis was performed in monkey DU, the only monkey with matched initial target locations recorded using a laminar probe. For each session, we identified the initial target location that yielded the highest decoding accuracy and compared it to the projected session location along the motion axis. Across sessions, maximum spike decoding accuracy showed a marginal correlation with projected location and tended to lie near the unity line (Supplementary figure 10L; Pearson correlation p = 0.062), whereas LFP decoding showed no significant relationship (Supplementary figure 10M). These results provide limited evidence that onset proximity influences velocity encoding strength.

Finally, we examined whether laminar depth influences velocity decodability. The SC exhibits a laminar organization in which neurons across depth contribute differentially to visual and motor processing, and afferent inputs are spatially distributed along the dorsoventral axis. Neurons in distinct layers may therefore differ in their contribution to velocity representation.

Because electrode penetrations were not perfectly aligned across sessions, we realigned depth profiles using the visual motor index (VMI, see Methods). After alignment, we evaluated velocity decodability at the single neuron level across layers. In this analysis, we were not concerned with temporal evolution of decoding accuracy, but rather with how strongly each neuron encoded velocity information across the entire analysis window. Thus, instead of reducing dimensionality across neurons, we applied PCA along the time axis, treating each neuron’s activity across time as a high dimensional vector (Figure 8A; see Methods). The resulting PCA scores were then used as input to LDA to classify the seven velocity conditions, following the same decoding framework used in the population analysis. This procedure yielded a single decoding accuracy value for each neuron. Across all three monkeys recorded with laminar probes, decoding accuracy peaked in dorsal intermediate layers or ventral superficial layers (Figure 8B-D); depth effects could not be assessed in the monkey recorded with the N-Form array. Compared with spike-based decoding, LFP-based decoding showed a flatter depth profile in one monkeys (DU), suggesting broader spatial integration of velocity related signals in LFPs.

**Figure 8.**
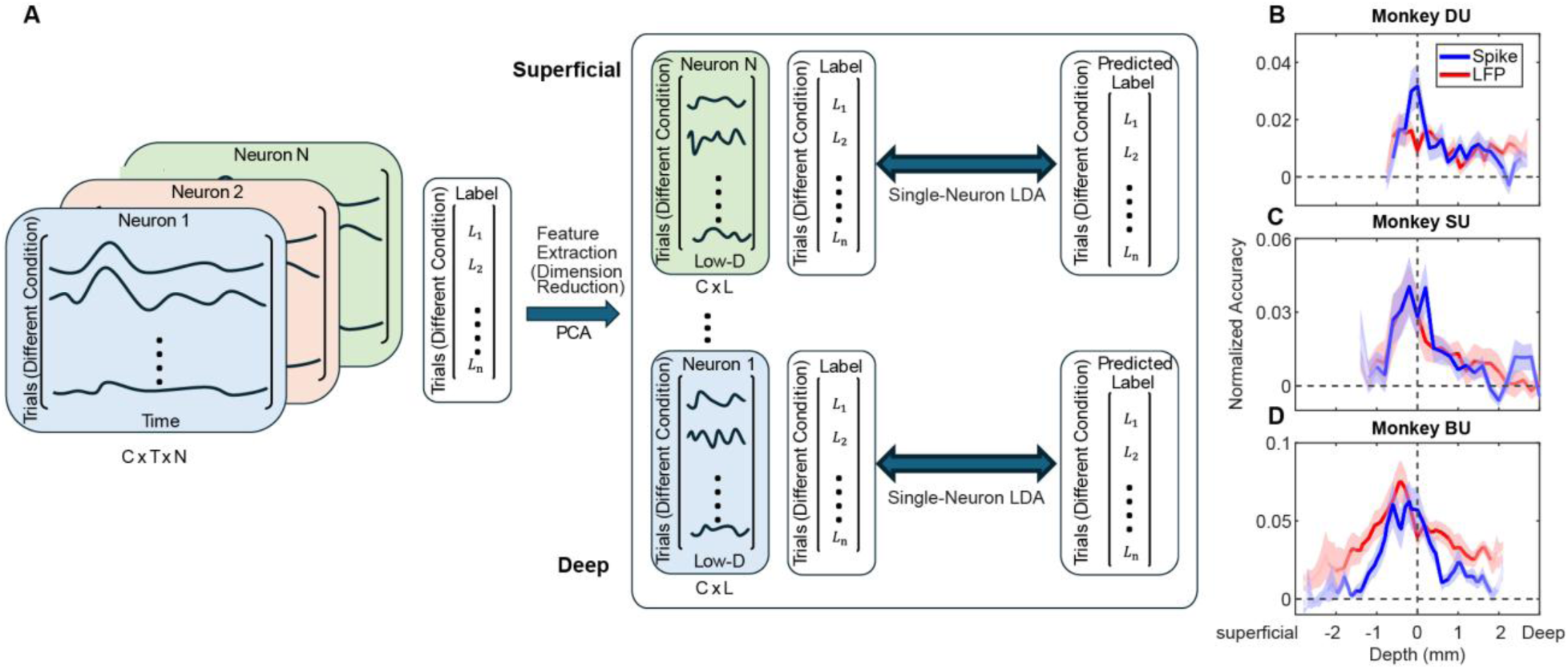
Illustration of dimensionality reduction method used for single neuron decoding analysis. **(A)** Spike density or LFP activity was first reduced along the temporal dimension with PCA to generate low-dimensional temporal features. Trials were labeled into seven speed conditions (outward 45°/s, 30°/s, 15°/s; inward 45°/s, 30°/s, 15°/s; stationary). Linear discriminant analysis (LDA) classifiers were trained separately on spike- and LFP-derived principal components. Decoding performance was evaluated using Monte Carlo cross-validation (80% training, 20% testing; 100 iterations). Decoding accuracy is plotted as a function of depth along the laminar probe for monkeys DU **(B)**, SU **(C)**, and BU **(D)**. Recording depth of 0mm on the x-axis denotes the transition between visual- and motor-dominant activity (VMI=0). The y-axis shows normalized decoding accuracy (actual minus shuffled), where 0 indicates chance level. Shaded regions indicate mean ± SE. The lighter trace at the extremes of the depth profile reflects estimates derived from fewer recording sessions.

Overall, anatomical position and motion direction did not show consistent relationships with decoding performance. In contrast, the relative spatial relationship between a neuron’s receptive field and the moving target trajectory showed a modest effect, with higher decoding accuracy observed when motion trajectory path closer to the receptive field center. Laminar depth exhibited the most prominent modulation, with decoding accuracy tending to peak in intermediate layers.

## DISCUSSION

Accurate orienting to moving objects requires the brain to compensate for the temporal mismatch between a target’s current position in the world and the delayed neural representation available to downstream motor circuits. The present findings identify the primate SC as a previously underappreciated locus of that computation. Its population activity carries robust information about target speed and motion direction in both spikes and LFPs, even though conventional single-neuron analyses revealed little consistent motion tuning. The combined use of dimensionality reduction and classification methods revealed that velocity is decodable shortly after visual onset, with decoding performance accumulating over time before reaching a plateau level. Moreover, we identified a V-shaped manifold in latent space, with the stationary condition at the vertex and speed-dependent trajectories diverging along separate arms for the two motion directions. Notably, this decodability cannot be explained solely by instantaneous target location. We also found that decoding performance does not systematically vary with factors such as rostrocaudal position or target motion direction. Instead, velocity related variance exhibits a pronounced laminar modulation, with intermediate layers showing higher decodability than superficial or deep layers. Together, these results redefine the classical view of the SC as a primarily spatial visuomotor structure, demonstrating that dynamic motion parameters are integrated into its population activity during visuomotor transformation ^44,50,56–58^.

### Traditional Single Neuron Analysis vs. Population Analysis

Traditional firing rate analyses emphasize mean rate differences across conditions and are well suited for identifying strongly tuned neurons. However, such approaches may underestimate distributed or temporally structured representations. Indeed, vector information is encoded in both spatial and temporal features of SC population activity ^59,60^. Recent studies across multiple brain regions have demonstrated that dimensionality reduction and population level analyses can reveal structured latent variables that are not apparent from single unit rate comparisons ^49,51^. Within the SC, ensemble approaches have identified representations beyond target location, including object related signals and decision related variables ^58,61–65^.

By applying dimensionality reduction to population activity combined with time resolved decoding, we detected coordinated patterns of activity across neurons and across time that are not readily captured by scalar firing rate comparisons alone. The persistence of above chance decoding after controlling for displacement differences further indicates that velocity related variance cannot be reduced to simple differences in mean firing rate attributable to target location. Decoding performance rose above chance at latencies consistent with known visual response delays in the SC and increased over time, suggesting progressive integration of motion related signals rather than a purely transient visual response.

Above chance decodability only indicates that information about target velocity is statistically present in the population activity and accessible to a linear classifier. However, decoding performance alone does not establish that this information is causally used by SC. Future studies combining perturbation, inactivation, or pathway specific manipulation with behavioral measurements will be necessary to determine whether and how SC activity contributes to the use of velocity information during interceptive saccades.

### Velocity Represented as a Unique V-shape Subspace

Low dimensional visualization revealed a structured organization of population activity associated with target speed and motion direction. In the reduced dimensional space, neural population corresponding to different target velocities formed a V shaped configuration that varied systematically with both speed and motion direction. This geometry was reproducible across sessions and was not attributable to displacement differences.

Analyses of the relationship between the target velocity subspace and the target location subspace revealed a consistent separation between the two representations across monkeys. Although the subspaces were not fully orthogonal, their relative orientations indicate that velocity information is organized largely independently of the dominant spatial representation. This conclusion is further supported by simulations showing that instantaneous target location alone cannot account for the observed decoding performance. Together, these results argue that SC population activity contains a distinct representation of target velocity rather than merely reflecting differences in target position. Some overlap between the two subspaces is nevertheless expected because velocity and displacement are inherently linked for moving stimuli. Thus, the observed geometry is most consistent with a representation in which motion information remains largely separable from, while still partially constrained by, the underlying spatial map.

These findings are consistent with the broader framework of representational subspace geometry observed in other brain regions, in which multiple task variables occupy different subspaces ^66–71^. Traditionally, the SC has been described as a retinotopic visuomotor map, with the/ dominant dimension of activity corresponding to target location and movement direction ^22,23,28–30^. Our results suggest that additional motion parameters, such as target velocity, are also embedded within the same population activity.

### Potential Sources of Velocity Information

A prominent finding of this study is the laminar dependence of velocity decodability. Neurons in the intermediate layers exhibited higher decoding accuracy than those in superficial or deep layers. This pattern aligns with prior reports showing that relative motion selectivity in the monkey SC is strongest in the deeper portion of the superficial layers, near the superficial-intermediate boundary ^46,47^. Together, these observations suggest that velocity information emerges preferentially at the interface between visual and intermediate layers. Multiple mechanisms could account for this laminar organization.

Intermediate-layer circuits may locally extract velocity information from visual inputs originating in the superficial layers. Anatomical and physiological studies indicate that intra-collicular connectivity differs across SC layers in many species ^72–74^, and neuronal cell types are heterogeneously distributed within the SC ^39,44,75,76^. Although their specific roles remain incompletely characterized in primates, work in mouse and hamster SC demonstrates that inhibitory microcircuits contribute to motion processing ^38,41^, raising the possibility that similar circuitry in macaque intermediate layers transforms visual motion signals into velocity-selective population codes. In this scenario, motion signals from retina and middle temporal (MT) area would first reach the superficial layers and then undergo local transformation within intermediate layers before being relayed to deep layers for motor execution. This is plausible given that MT is a major source of cortical speed and direction information ^77–79^, although the precise role of MT projections to the superficial layers of SC remain unclear. Visual response latencies in area MT can range from 35ms to 325ms, depending on stimulus type ^80,81^, which is comparable to or later than the onset of SC visual responses depending on monkeys (supplementary figure 4E). A retinal contribution could also account for the early emergence of velocity information in the SC. The earliest decodable velocity signals in SC are unlikely to depend exclusively on cortical input and may instead reflect rapid feedforward retinal drive and early intra-collicular processing. Consistent with this view, early velocity decodability was present but weak, suggesting an initial, coarse visual representation that is subsequently refined, potentially through recurrent intra-collicular circuits and later-arriving cortical inputs.

Velocity-related signals may arrive directly at intermediate layers from cortical areas such as the frontal eye field (FEF), which projects to these layers and carries motion-related information ^82,83^. Additionally, studies in cats show that different types of retinal ganglion cells project selectively to distinct SC layers ^40^, raising the possibility that velocity-sensitive retinal inputs preferentially target the lower superficial layers in monkeys as well.

Most plausibly, both mechanisms operate together. Early, relatively coarse velocity information may arise from retinal and superficial-layer drive and local intermediate-layer processing, accounting for the initial above-chance decoding. As cortical motion areas such as MT and FEF become engaged, additional velocity-selective signals may converge directly onto intermediate layers, strengthening and stabilizing the population representation. This layered convergence model explains both the laminar peak in decoding accuracy and the temporal evolution of velocity decodability.

Future pathway-specific perturbations combined with simultaneous population recordings across MT, FEF, and SC will be necessary to dissociate inherited versus locally generated velocity signals and to determine how laminar circuits shape the geometry of velocity representations in the SC population code.

### Local Field Potentials vs. Spikes

Spikes and LFPs reflect different aspects of neural activity. Spikes represent high-frequency, thresholded action potentials from neurons near the electrode tip, whereas LFPs are obtained via low-pass filtering and reflect slower fluctuations in extracellular potential. Although these signals are correlated, numerous studies indicate that LFPs primarily reflect synaptic input and coordinated population activity within a local network rather than the output of individual neurons ^84–86^. Consistent with this, LFPs typically exhibit greater similarity and lower spatial variability across nearby recording sites compared to spikes.

We therefore asked whether both spikes and LFPs encode target velocity, and whether they do so in a similar manner. Overall, the results were remarkably consistent across signal types. For both spikes and LFPs, velocity was decodable above chance shortly after target onset, with comparable onset latencies of decoding. In addition, low-dimensional projections revealed similar V-shaped subspace geometries for velocity representation in both signals.

The primary difference emerged in laminar analyses. In one of the three monkeys, LFP decoding accuracy exhibited a flatter profile across depth compared to spikes (Figure 8B). Furthermore, in the one monkey (same monkey in Figure 8B) for which we could examine the relationship between recording site and the target initial location associated with maximal decoding accuracy, no systematic relationship was observed for LFPs (Supplementary Figure 10L). These findings are consistent with the spatial integration properties of LFPs. Since they reflect pooled local population activity and synaptic inputs over a broader spatial extent, they are expected to exhibit less laminar specificity and less site-to-site variability than spiking activity recorded from the same electrode.

To test whether spikes and LFPs carried complementary information, we also tried combining their PCA scores and repeating the decoding analysis. This joint analysis did not yield consistent significantly improved decoding performance across sessions and monkeys, making it hard to interpret whether both signals have largely overlapped velocity-related variance in the recorded population.

Further work is needed to disentangle the distinct functional roles of spikes and LFPs in velocity representation within the SC. Future analyses could include frequency-resolved decoding and current source density analysis, which may reveal whether specific LFP frequency bands or laminar or spatial input patterns differentially contribute to motion processing. Such approaches may clarify whether spikes and LFPs represent distinct computational stages of velocity encoding or instead reflect a shared underlying population dynamic.

## METHODS

### Animal Preparation

All experimental and surgical procedures were approved by the Institutional Animal Care and Use Committee of the University of Pittsburgh and were in accordance with the guidelines of the U.S. Public Health Service policy on the humane care and use of laboratory animals. Four male rhesus macaques (Macaca mulatta) participated in this study. Surgeries were performed under sterile conditions with isoflurane anesthesia. The procedures for monkeys DU, SU, and BU have been reported in earlier studies ^87,88^. Briefly, a recording chamber was implanted at a 40° posterior tilt in the sagittal plane and centered on the midline to provide orthogonal access to both left and right SCs (Figure 1A). When required, a 3D-printed angular adaptor was used to fine-tune the penetration angle. For monkey LO, a custom-designed, N-Form multi-electrode array (Plexon, Inc and Modular Bionics, Inc) was implanted in the SC to cover a 1.2mm × 1.2mm region of the SC. Access to the SC also followed a posterior approach. A D-shaped flap of the skull was removed over the posterior parietal and extrastriate cortices, followed by a durotomy of expose the medial wall of the underlying cortex. The two cortical hemispheres were pushed apart by a few millimeters. A small portion of the exposed posterior corpus callosum was cut, the blood vessels separated, and the pineal gland removed to expose the dorsal aspect of the SC. The N-Form array was then implanted in the surface of the right SC (Figure 1B). The probe cable traveled along the surgical approach penetration to the cortical surface, where a custom-designed 90-degree bend in the cable allowed it to turn and coarse anteriorly. The Omnetics connectors at the end of the cable were placed in a custom-designed titanium block and implanted on the skull over the frontal bone. The dura was sutured closed and the D-shaped bone flap was placed back in place and attached to the skull. Following surgery, the monkey was returned to its home cage for full recovery, with postoperative administration of antibiotics and analgesics according to protocol.

We disclose that monkey DU initially underwent an N-Form array surgery, but the neural signal was not reflective of the SC. We therefore performed the standard chamber placement surgery explained above at a later date to permit acute recording experiments.

### Experimental Setup

Animals were trained on behavioral tasks prior to surgery. Visual stimuli presentation, behavioral control, and data acquisition were handled using custom software: a LabVIEW-based interface for monkeys SU and BU ^89^, and a Psychtoolbox-based interface ^90–92^ for monkeys DU and LO. An LED-backlit flat-screen monitor (120 Hz, 1080p) was used to display white pixels (0.5° × 0.5° or 0.2° × 0.2° visual angle) against a dark gray background. To synchronize timing across experimental stages, a white square was simultaneously flashed at the screen corner and recorded via a photodiode. This region of the monitor was covered so that the white square and photodiode were not visible to the animal. Head restraint was maintained using a thermoplastic mask attached to the primate chair ^93^. Eye position was sampled at 1 kHz with an infrared eye tracker (EyeLink 1000, SR Research) and calibrated at the start of each session. Saccade onset and offset were identified using standard velocity thresholds. Monkeys received liquid rewards controlled by a computer-operated solenoid valve.

### Neurophysiological Recordings and Microstimulation

Neural recordings in the SC were performed using Alpha Omega (monkey BU, 24 channels, 150μm intercontact distance, 264μm diameter) and Plexon S-probes (monkeys BU, SU, and DU; 24 channels; 210µm diameter; 200µm intercontact spacing in monkeys BU and SU, 150µm in monkey DU), and the N-Form Array (monkey LO; partnership between Plexon Inc and Modular Bionics Inc; 128 channels, 16 shanks in 4×4 arrangement, 400μm distance between shanks, 250μm intercontact distance, 125μm shank diameter). In all configurations, the electrodes tracked a path approximately orthogonal to the SC, traversing its dorsoventral axis within a column to encounter neurons with similar optimal visual and motor vectors.

The SC was identified based on neural responses (“swishes”) to visual stimuli and/or before saccades, or stimulation-evoked saccades ^23,26,94^. Raw neural signals were sampled at 30kHz using the Scout system (Ripple Neuro LLC), visualized with Trellis software (Ripple Neuro LLC), and integrated into the central, custom-built data acquisition system. The neural signals were separated into spikes and LFPs, the latter downsampled to 1 kHz. The laminar probe was positioned to ensure that recording contacts covered a substantial portion of the SC along the dorsoventral axis, spanning superficial to deeper layers.

Spikes were detected using a threshold-crossing criterion and isolated with a spike-sorting algorithm (MATLAB based MLIB - toolbox). For each recording contact, we selected at most one neuron per site, defined as the unit with the strongest firing rate and clear tuning to the visual stimulus. If a well isolated neuron was not identified, we retained the noisy signal to preserve a consistent data structure across sessions for subsequent population analyses. Spike trains were visualized as raster plots and analyzed using spike density functions (Gaussian kernel, σ=6) ^95^.

Although precise neuronal location was not critical for this study, we estimated recording locations within the SC topographic map using stimulation evoked saccade amplitudes in monkeys DU, SU, and BU (Figure 1E, H, & I). Microstimulation (40μA, 400Hz, 200ms; biphasic pulses with 200μs phase duration and 17μs interphase interval) was applied at the beginning and end of each session to localize recording sites. For each channel, three stimulation trials were performed, and the mean evoked saccade amplitude was used to define the neuronal preferred location. Visual field locations were converted to SC map coordinates using the logarithmic mapping function of Van Gisbergen and colleagues ^96,97^. For monkey LO, microstimulation was not performed. Instead, neuronal locations were estimated based on the target position that elicited the strongest visual response to a stationary dot stimulus. In addition, given the fixed geometry of the N-form array, array placement was estimated by fitting estimated neuronal locations across sessions using a least squares procedure (Figure 1F, Supplementary Figure 1). Recording stability was assessed based on consistency of spike sorting results and the stability of neuronal preferred visual locations across sessions.

### Behavioral Task

Monkeys were trained using operant conditioning to perform several randomly interleaved oculomotor tasks. The “Delayed Stationary Target Task” and “Delayed Moving Target Task” required generating eye movements to stationary or moving targets, respectively, whereas the “No-Go” condition required maintaining central fixation despite the presence of a peripheral stimulus of either type (Figure 1C). All four trial configurations were randomly interleaved.

Each trial began with the animal fixating a central stimulus. After a brief fixation period, another target appeared in the visual field, but the animal had to maintain gaze on the fixation point. In the Delay Saccade Task, the target remained stationary, while in the Delayed Moving Target Task, it moved immediately upon onset. On saccade trials, the disappearance of the fixation point a short period of time (150-500ms for monkey DU and LO; 50-500ms for Monkey SU and BU) after target onset served as a “go cue” for the animal to initiate a saccade toward the target. Following the saccade, the monkey was required to continue fixating the stationary stimulus or tracking the moving target for 250-500ms to receive a liquid reward. In the No-Go task, the fixation point remained illuminated (400 to 1000ms) for the remainder of the trial.

The target’s direction of motion was tailored to the neural recordings and differed across animals. For monkey DU, targets moved along the direction axis preferred by most neurons at the recording site (Figure 1E), whereas for monkey LO, target motion axis covered the range of the recorded population (Figure 1F). For Monkeys SU and BU, the moving target was confined to the horizontal axis (Figure 1H-I).

In motion trials, the target moved either toward (“inward” trials) or away from (“outward” trials) central fixation at one of three speeds: 15°/s, 30°/s, or 45°/s. For Monkeys DU and LO, initial target locations were matched across target velocities (Figure 1D). For Monkeys SU and BU, initial locations are varied among different velocities (Figure 1G). Initial target position also varied between Go and No-Go tasks (Supplementary Table 1), in part due to limits of the oculomotor range and the spatial sensitivity of the eye tracker. Note that positive target location values correspond to the contralateral visual hemifield relative to the recording side.

### Data Analysis

All data analyses were conducted using custom code written in MATLAB (MathWorks, Inc.). Visual responses were examined at both single-neuron and population levels. Analyses included both Go and No-Go trials and considered both successful trials and failures in which the monkey did not complete interception or post-saccadic pursuit. This decision was deemed acceptable since we focused on visual and delay period activity that preceded saccade onset by at least 100 msec. The duration of the visual analysis window was matched across all target motion conditions within each monkey (300ms for monkeys DU and LO; 200ms for monkeys SU and BU). One reason for this limit was the shorter delay duration needed to account for fast speed trials in the Go condition trials. To isolate visually driven activity and retain a sufficient number of trials for subsequent analyses, we excluded trials in which eye speed exceeded 80°/s during the visual period or within 100ms after the end of the visual window. This threshold removes large eye movements but does not eliminate microsaccades or slow eye drift. We therefore quantified eye drift separately and concluded that it occurred too late to influence our findings (Supplementary figure 5). Baseline activity was defined as the time preceding target onset from the same set of trials.

### Realignment of Neuronal Depth Profiles across Sessions

Neuronal depth profiles were realigned across sessions using the visual motor index, defined as 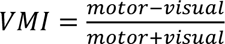, where visual activity was computed as the average firing rate within a ±10ms window around the visual response peak, and motor activity was computed as the average firing rate within a ±10ms window around the motor response peak. Following realignment, channel zero (VMI = 0) corresponded to the transition from visually dominated to motor dominated activity.

For sessions in which no clear transition from visually dominated to motor dominated activity was observed, or in which some channels lacked both visual and motor responses, the transition layer was defined as the layer between the visual and motor response peaks.

### Principal Component Analysis

PCA reduces the dimensionality of the observed data by identifying orthogonal axes that capture the greatest variance while minimizing the impact of noise ^53,54^. In our initial analysis, we applied PCA to each session’s data across neurons, while preserving the temporal structure within a trial (Figure 3). This analysis was applied separately to spike density traces and LFP waveforms.

Before dimensionality reduction, activity from each trial was normalized by subtracting the mean and dividing by the standard deviation, yielding zero mean and unit variance. The normalized trials were then concatenated across conditions and time points to form the population activity matrix used for decomposition.

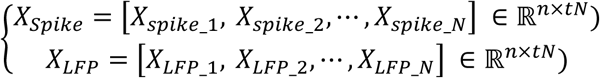

where *X_N_* ∈ ℝ*^n^*^×*t*^, n is the number of neurons, t is the number of time points during the baseline (50ms) and visual period (300ms for monkeys DU and LO; 200ms for monkeys SU and BU), and N is the total number of trials. Singular Value Decomposition (SVD) was used to perform the PCA:

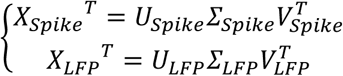

where

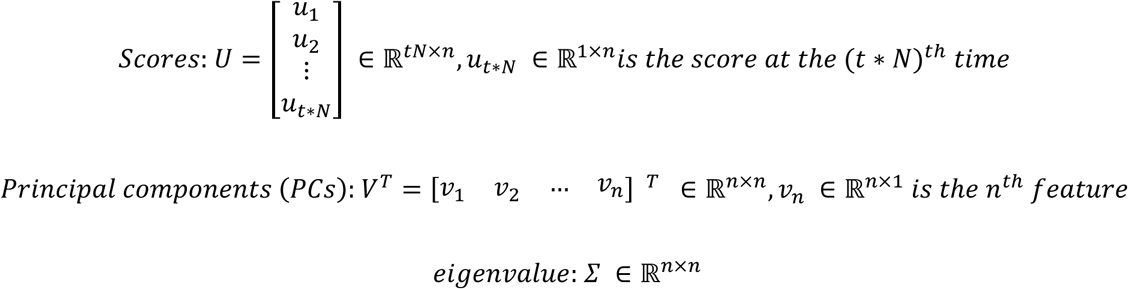

After selecting the first *i* dimensions, the reduced score matrix is:

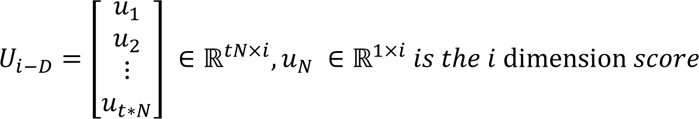

After obtaining the low-dimensional concatenated scores, we separated them back into individual trials:

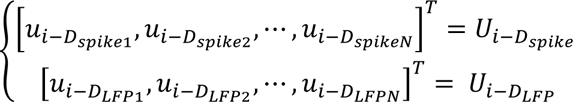

where *u_i_*_−*D*_ ∈ ℝ*^t^*^×*i*^ is the time-varying low-dimensional representation of each trial.

In another analysis, we performed PCA on the activity of individual neurons. In this case, PCA was applied along the time axis of each trial (Figure 8A), separately for spikes and LFPs, yielding a low-dimensional representation of temporal structure for single-neuron activity. PCA were performed separately for each neuron’s baseline (200 to 0ms before target onset) and visual period (0 to 180ms after target onset). Before decomposition, each trial was normalized to zero mean, and then all trials were concatenated differently from previous PCA:

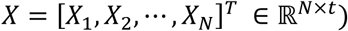

where *X_N_* ∈ ℝ*^t^*^×1^ is the neural activity for each neuron, and t is the number of time points during the baseline or visual period. SVD was used to compute the PCA:

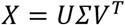

After selecting the first i PCs, the reduced score matrix was:

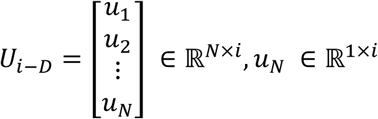

where *u_N_* represents the low-dimensional projection (scores) of the *N*^th^trial.

### Linear Discriminant Analysis

For each trial, we created a label vector: 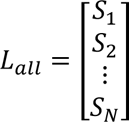, where *S*_N_ is the velocity category of the *N^t^*^ℎ^ trial. For the binary LDA analysis, the category indicated either a stationary or moving target condition. For the multi-class LDA analysis, outward motions were assigned positive labels, inward motions negative labels, and stationary targets were labeled as 0. In addition, each unique velocity value was treated as a separate class (Figure 3, ①). At each time point *t* of the score matrix u, we averaged across a sliding window ([-10ms,10ms]) to obtain the mean score vector *u_i_*_−*D*_(*t*) ∈ ℝ^1×*i*^ at time *t*. We then constructed both types of LDA decoders:

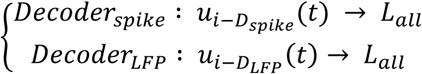

where *u_i_*_−*D_spike_*_ and *u_i_*_−*D_LFP_*_ are the low-D PCA score matrices obtained across neurons for population-level analysis. Cross-validation based regularized LDA (MATLAB function cvshrink, https://www.mathworks.com/help/stats/classificationdiscriminant.cvshrink.html) was used to avoid overfitting.

Similarly, for single-neuron analyses, we used PCA scores obtained by performing dimensionality reduction along the temporal axis. LDA decoders were trained separately for each neuron using its corresponding low-dimensional score matrix *U_i_*_−*D*_ and decoding was performed across all velocity labels (*Decoder*: *U_i_*_−*D*_ → *L_all_*; Figure 8A).

### Cross-Validation

Monte Carlo cross-validation ^98^ was used to evaluate decoder performance. For each session, trials were stratified by stimulus condition (across initial location × velocity category for monkey DU and LO; across velocity category for Monkey SU and BU) to ensure balanced representation in both training and test sets. In each iteration, 80% of trials were randomly assigned to the training set and the remaining 20% to the testing set. LDA parameters were estimated from the training set and then applied to the test set without re-fitting. Decoder accuracy was computed as the proportion of correctly classified test trials. This process was repeated for 100 iterations, yielding a distribution of decoding accuracies.

### Visualization - Velocity Subspace

Because LDA provides a linear transformation that maximizes category separability, we used its components to visualize neural activity in a low-dimensional subspace (Figure 3,②). Since LDA was applied independently at each time point, each time step yields a different transformation that optimally separates categories at that moment. To examine how population activity evolves within a consistent subspace, we fixed the transformation matrix at a single reference time window (180 ± 10ms after target onset) and projected activity from all time points into that same subspace.

At 180ms, the LDA transformation matrices were obtained as:

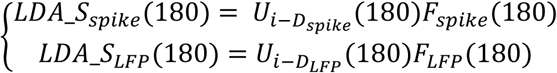

where *LDA*_*S* denotes the low-dimensional score obtained by projecting neural activity onto the LDA discriminant components. We then projected scores from all other time points *t* into this fixed subspace using this transform matrix *F*(180):

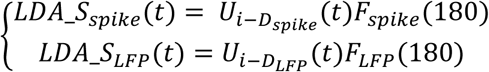

For visualization, we projected population activity onto the first two LDA components and plotted the resulting low-dimensional representations over time to illustrate the evolution of population trajectories. To enable cross-session and cross-monkey comparisons of the low-dimensional subspace geometry, we applied normalization and alignment steps at the same reference time window (180 ± 10ms after target onset) as the transformation matrix, and the resulting normalization matrix was applied consistently to all other time points.

First, the stationary target subspace was centered at the origin. Specifically, the centroid of the stationary and moving category was translated to:

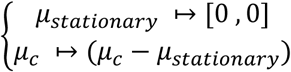

where *μ_c_* is the center of subspace of category c (e.g., outward motion at 45°/s).

Second, the relative locations of moving target categories were rescaled with respect to the stationary subspace. For each session, category centroids were normalized such that the maximum value across categories was set to +1 and the minimum to −1:

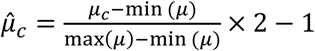

where *µ^_c_* denotes the rescaled centroid.

Third, axes were flipped when necessary to ensure a consistent orientation across sessions, such that outward motion was located on the left and inward motion on the right.

Finally, to align category structure across sessions, subspaces were rotated to minimize the following loss function:

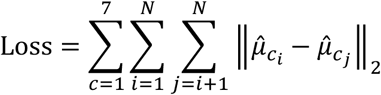

where *c* indexes the seven velocity categories, *N* is the total number of sessions, and *µ^_ci_* is the centroid of category *c* in session *i*. We then plotted the trajectories of population activity in this subspace over time, grouping trials by velocity category.

### Visualization - Location Subspace

We also projected the population activity onto the location subspace. The transformation matrix was obtained via linear regression, projecting the neural population scores *U_i_*_−*D*_ onto the target location label *L* at 70ms after target onset (roughly corresponding to the visual burst that encodes the target’s spatial location):

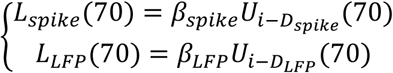

where *β* represents the regression coefficients. We then applied *β* across all other time points to obtain the predicted location labels dynamically:

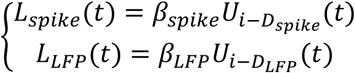

We then visualized the population trajectories in the location subspace over time. Trials were also grouped by velocity category to assess whether velocity-dependent differences were already separable within this spatial representation.

### Simulations

To determine whether target displacement alone could account for the observed velocity decodability, we simulated neural population activity based solely on target location. Forty simulated neurons were distributed uniformly across a 4 mm region of the SC map, spanning uniform locations from the foveal (0 mm) to peripheral (4 mm) representations. Each neuron was assigned a log-Gaussian receptive field in visual space - equivalently, a Gaussian point-image in SC space centered at its location with a standard deviation of 0.2 mm (Figure 6A). To introduce trial to trial variability, uniformly distributed noise with an amplitude of ±10% of the neuron’s peak response was added independently to all neuronal responses on each simulation iteration (Figure 6B). In the absence of prior systematic characterization of SC receptive fields during moving-target conditions, we assumed that receptive fields during motion follow the same log-Gaussian profile as those measured for stationary targets. Although increased intro-collicular neuronal interactions are expected immediately after target onset for both stationary and moving targets, potentially affecting overall firing rate amplitude, we assume the relative tuning across target locations remains stable. As long as this relative structure is preserved, separability of target locations in population activity should not be affected. To simplify the simulation, we therefore assumed identical receptive field amplitudes for stationary and moving targets over time.

Population responses were generated for target locations spanning −10° to 35°, encompassing the full range of locations traversed by the moving targets. A linear regression model similar as the location axis was trained to estimate target location from the simulated population activity (Figure 6C). We first project the simulated neural population activity *P* onto the target location label *L* at 70ms after target onset (roughly corresponding to the visual burst that encodes the target’s spatial location):

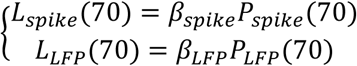

where *β* represents the regression coefficients. The trained decoder *β* was subsequently applied to all time points without retraining:

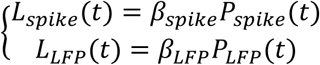

Using the known target trajectories for the matched and unmatched initial location conditions (Figure 6D & F), we generated decoded location trajectories for the matched and unmatched conditions.

For the matched initial location condition, 30 trials were simulated for each combination of target velocity and initial location, matching the distribution of trials in the experimental data. For the unmatched initial location condition, 120 trials were simulated for each velocity category. These trials were divided equally among all initial target locations associated with that velocity category, such that each initial location contributed the same number of simulated trials. The resulting decoded location estimates were then analyzed using the same binary and multiclass LDA procedures applied to the experimental data, allowing us to quantify the extent to which target displacement alone could account for the observed velocity decoding performance.

We also repeated the analysis in a smaller neuronal population matching the spatial scale of our recordings. Ten simulated neurons were uniformly distributed within a 1 mm region of the SC map, centered at rostral (1 mm), middle (2 mm), and caudal (3 mm) locations. All other analyses were identical to those described above.

### Statistics

Statistical significance across time series data was assessed using a cluster-based permutation test introduced by Maris and Oostenveld ^99^ and implemented with the MATLAB function permutest (https://www.mathworks.com/matlabcentral/fileexchange/71737-permutest). At each time point, decoding accuracy was compared against chance level and was grouped into clusters. Statistical significance was determined by comparing each cluster’s statistics to a null distribution generated through 10000 random permutations. This approach controls for multiple comparisons across time while accounting for temporal dependence in the data. For visualization, significant time periods indicated by horizontal lines, with only the cluster of time exhibiting the lowest p-value shown for each combination of decoder type and velocity category.

For analyses of noncontiguous data, statistical comparisons between groups were performed using either independent or paired t-tests, as appropriate. Multiple comparisons were controlled using the Benjamini Hochberg procedure. Pearson correlation analysis was used to assess the relationship between pairs of variables.

To estimate chance level performance for statistical comparisons, we repeated the decoding procedure after randomly shuffling trial labels within each session. This preserved the structure of the neural data while disrupting the relationship between neural activity and target velocity. The resulting distribution of decoding accuracy served as a null distribution, against which the observed decoder performance was evaluated.

## ACKNOWLEDGEMENTS

We thank Drs. J. P. Herman, C. Huang, and J. P. Mayo for scientific discussions and critical feedback on previous versions of the manuscript; Dr. R. C. Saunders for contributions to the N-form array surgery; Dr. K. Mohsenian for participation of data collection; Drs. A. Fisher and V. Mrotz, surgical assistants S. L. Cashman and N. E. McLane, and the veterinary technicians and husbandry staff of the Division of Laboratory Animal Resources for assistance with animal care; and Dr. I. Smalianchuk for programming assistance. The study was funded by the following NIH grants: R01EY037837 and P30EY008098.

## AUTHOR CONTRIBUTIONS

F.Y. and N.J.G. conceived and designed the study. F.Y. performed the experiments, analyzed the data, and performed the simulations. C.B. contributed to data collection and edited the manuscript. M.A.G.E. performed the N-form array surgery. F.Y. and N.J.G wrote the manuscript.

## CODE & DATA AVAILABILITY

Source data and custom MATLAB codes that support the findings of this study are available from the corresponding author upon reasonable request.

## MATERIALS AND CORRESPONDENCE

Correspondence and requests for more information should be addressed to N.J.G.

