## Supplemental Figures and Table for "Distinct neural geometries for target position and velocity in the primate superior colliculus"

| | Electrode | Speed | Initial Location (Go Task) | | Initial Location (No-Go Task) | |
|---|---|---|---|---|---|---|
| | | | In | Out | In | Out |
| Monkey DU | Laminar | 15 | 15,20 | 5,10 | 5,10,15,20 | 5,10,15,20 |
| | | 30 | 15,20 | 5,10 | 5,10,15,20 | 5,10,15,20 |
| | | 45 | 15,20 | 5,10 | 5,10,15,20 | 5,10,15,20 |
| | | Stationary | 5,10,15,20 | | 5,10,15,20 | |
| Monkey LO | N-Form Array | 15 | 15,20 | 5,10 | 5,10,15,20 | 5,10,15,20 |
| | | 30 | 15,20 | 5,10 | 5,10,15,20 | 5,10,15,20 |
| | | 45 | 15,20 | 5,10 | 5,10,15,20 | 5,10,15,20 |
| | | Stationary | 5,10,15,20 | | 5,10,15,20 | |
| Monkey SU | Laminar | 15 | 10,16,22,26 | 2,7,12 | 26 | 2 |
| | | 30 | 14,18,23,26 | 2,4,10 | 26 | 2 |
| | | 45 | 17,21,26 | -4,2,6 | 26 | 2 |
| | | Stationary | 4~24 | | Not performed | |
| Monkey BU | Laminar | 15 | 10,16,22,26 | 2,7,12 | 26 | 2 |
| | | 30 | 14,18,23,26 | 2,4,10 | 26 | 2 |
| | | 45 | 17,21,26 | -4,2,6 | 26 | 2 |
| | | Stationary | 4~24 | | Not performed | |

**Supplementary Table 1.** The chart shows the distribution of initial target position according to animal, target condition (stationary or moving), task type (go or no-go), and motion direction (inward or outward). Note that initial target position and speed were matched for monkeys DU and LO but unmatched for monkeys SU and BU. Also, data were recorded with a N-form array in monkey LO and a multicontact laminar probe in the other three animals.

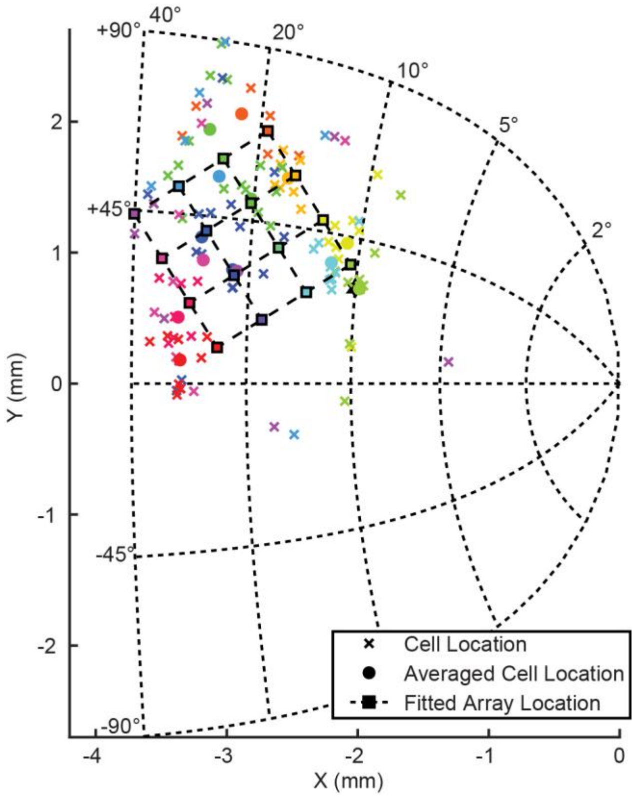


**Supplementary Figure 1.** Estimated recording locations for monkey LO using the N-Form array geometry. Each “x” marker represents the estimated SC location associated with an individual electrode contact from a recording session. Each location is determined by transforming the stimulation-evoked vector into SC space. Markers with the same color correspond to the same electrode contact across sessions. Circle markers indicate the average estimated location for each recording site across sessions. Based on these average locations and the geometry of the N-Form array, we fitted a 4 × 4 array configuration using a least squares procedure. Dashed grid lines illustrate the fitted array location, and square markers with the same color denote the corresponding fitted recording locations for each electrode contact.

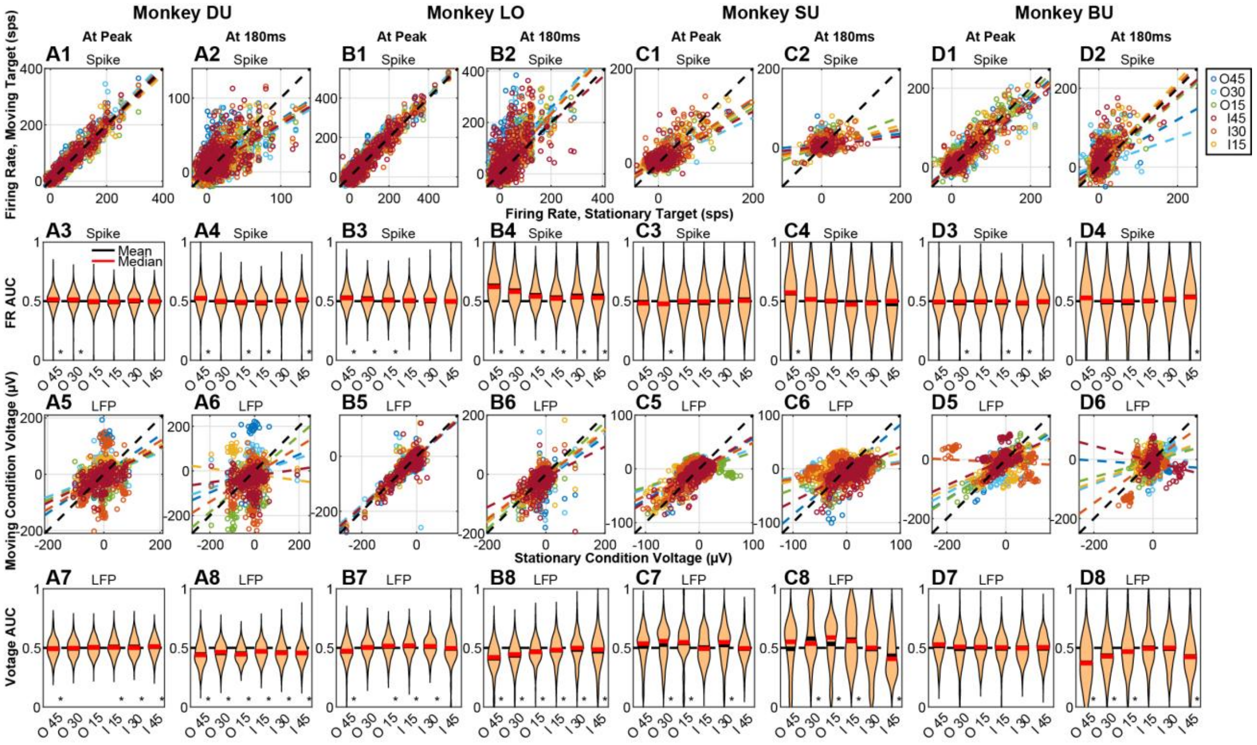


**Supplementary Figure 2.** Comparison of activities of individual neurons during moving and stationary target conditions. **(1 & 2)** Scatter plot of peak firing rates during the initial visual response **(1)** and activity at 180ms after target onset **(2)** for moving targets versus their matched stationary conditions for each monkey (**A**, DU; **B**, LO; **C**, SU; **D**, BU). Each point represents a single moving target condition compared to its stationary condition matched for the same initial position. Thus, data from an individual neuron occupy six points. Colored dashed lines denote linear fits for each speed condition. **(3 & 4)** Violin plots of the area under the receiver operating characteristic curve (AUC) computed from single-trial peak firing rates **(3)** and firing rates at 180ms **(4)**, quantifying discrimination between moving and stationary conditions for each speed. Data are matched for initial target location and then averaged across speed conditions and sessions for each monkey. Black and red lines respectively denote mean and median AUC values. Asterisks denote statistical significance (Benjamini-Hochberg correct t-test, $p<0.05$). **(5 & 6)** Same analysis as in **(1 & 2)**, applied to LFP trough amplitudes **(5)** and LFP activity at 180ms after target onset **(6)**. **(7 & 8)** The ROC analysis is applied to LFP trough amplitudes **(7)** and LFP activity at 180ms after target onset **(8)**. Same configuration as **(3 & 4)**.

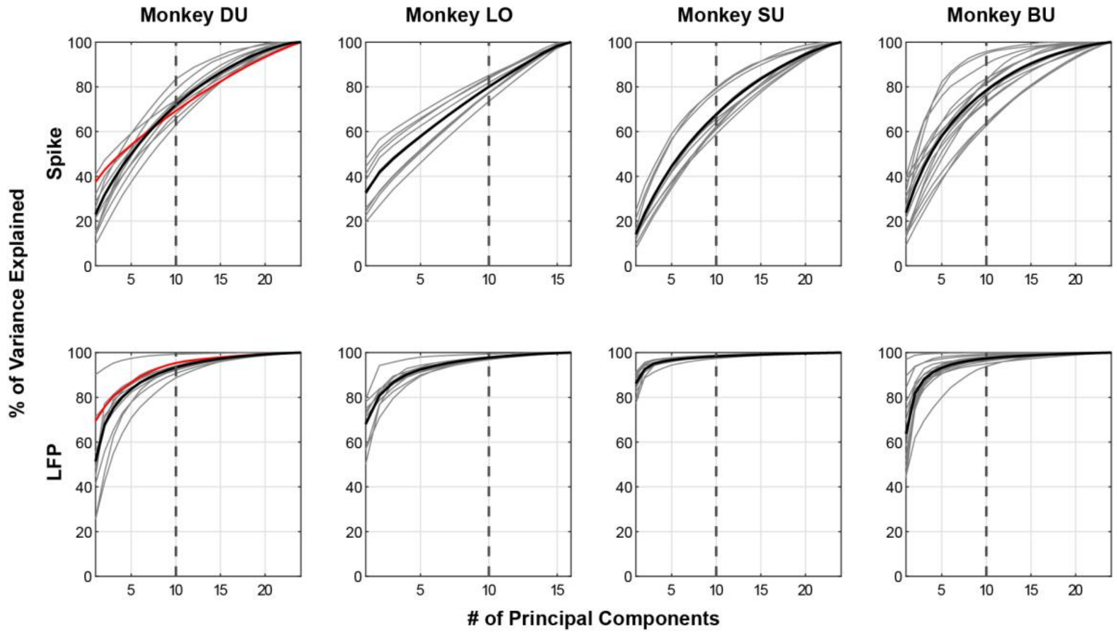


**Supplementary Figure 3.** PCA performance. Each panel plots the cumulative percent variance explained as a function of number of principal components. Each thin trace refers to an individual recording session. The red trace denotes the example session illustrated in Figures 2 and 4. The thick black traces is the average across all sessions. The top and bottom rows plot this metric for spike and LFP data, respectively. Each column shows data from one animal. The vertical dashed line indicates the number of PCs retained for subsequent analyses.

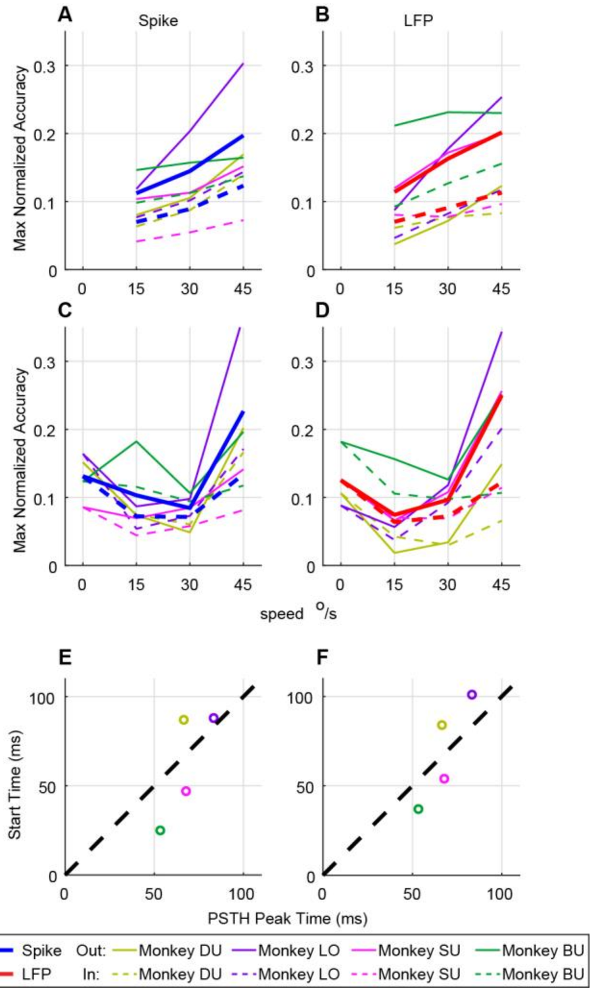


**Supplementary Figure 4.** Normalized overall decoding accuracy as a function of target speed. **(A & B)** Maximum normalized decoding accuracy across the entire analysis window (0 to 300 ms after target onset for DU and LO; 0 to 200 ms after target onset for BU and SU) for the binary decoder is plotted against target speed for both spike and LFP signals. Solid and dashed lines represent outward and inward motion, respectively. The color of each thin trace indicates animal identity. Each thick trace is the average of thin traces for each motion direction. **(C & D)** Maximum normalized decoding accuracy across the same analysis window for the seven-category decoder for spike and LFP signals. Same configuration as top row. **(E & F)** Relationship between the onset time of statistically above-chance decoding accuracy and the spiking activity peak time for spike and LFP decoders. Spike peak time was computed for each session based on the neuron with the highest firing rate at its preferred location and then averaged across sessions within each monkey. The same spike peak time measure was also used for comparisons with the LFP decoder, since a clear LFP trough was not consistently observed across sessions. Each point represents one monkey. The dashed diagonal line indicates unity slope.

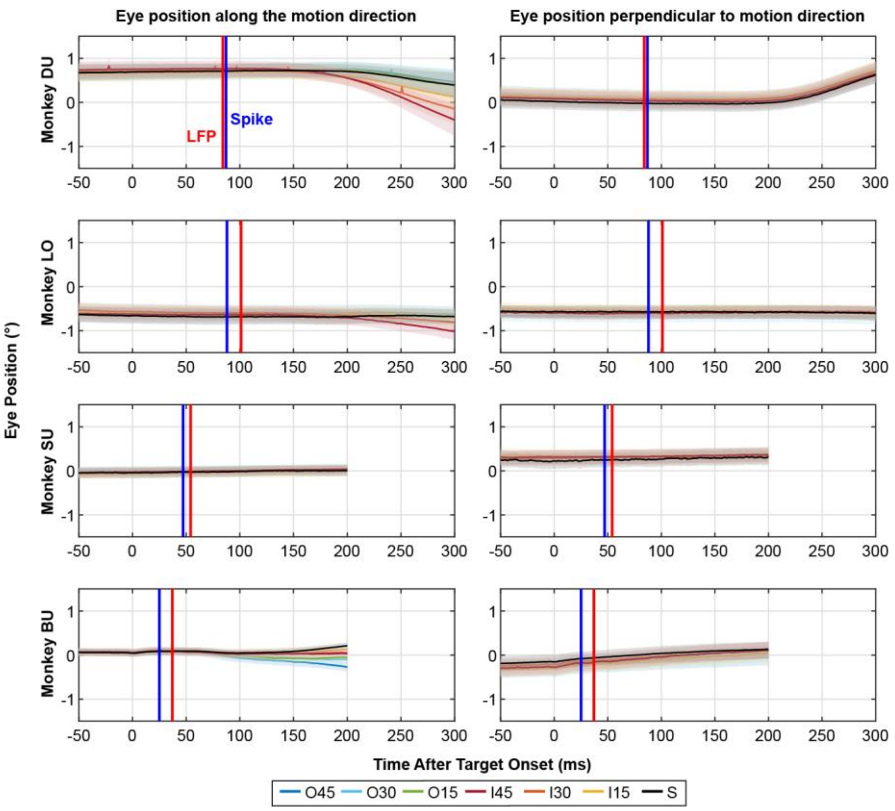


**Supplementary Figure 5.** Eye drift across target speeds and motion directions. Each row plots eye position traces aligned to target onset for one monkey. The left column shows eye position projected along the motion axis, while the right column shows the orthogonal component. Colored traces represent different target speed conditions, with shading indicating variability across trials (mean ± SE). Vertical lines mark the time at which significant speed decoding emerges from spike activity (blue) and LFP signals (red), as determined by cluster-based permutation tests in Figure 4. Key observation is that speed decoding occurs before and therefore is not a result of drifts in eye position.

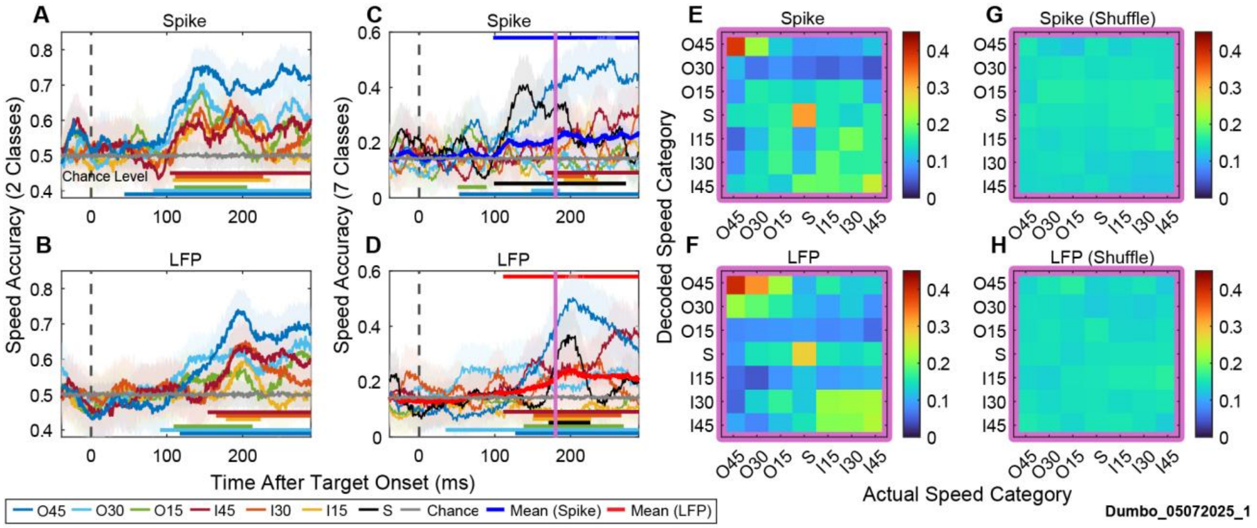


**Supplementary Figure 6.** Time-resolved decoding of target speed for an example recording session (Same session as in Figure 2). **(A & B)** Binary decoding accuracy separating each moving speed condition from the stationary condition (2-class classification) for spike and LFP activity, respectively, is plotted as a function of time. Target onset is indicated by the dashed vertical line. Shaded regions show mean ± sd across cross-validation iterations. Gray traces denote chance level. Colored horizontal bars at the bottom of each panel mark the contiguous time clusters with decoding accuracy significantly above chance exhibiting the lowest p-value (cluster-based permutation test). **(C & D)** Multiclass decoding accuracy for discriminating all seven speed conditions (outward 45°/s, 30°/s, 15°/s; inward 45°/s, 30°/s, 15°/s; stationary) for spike and LFP activity, respectively, is plotted as a function of time. Conventions are as in **(A & B)**. Colored horizontal bars at the top of each panel mark the contiguous time clusters with decoding accuracy significantly above chance exhibiting the lowest p-value (cluster-based permutation test) for the averaged across categories decoding accuracy. The vertical purple line indicates the time point selected for confusion analysis. **(E & F)** Confusion matrices show the proportion of trials decoded as each speed category for spike and LFP activity. Rows correspond to decoded speed and columns to actual speed. **(G & H)** Corresponding confusion matrices obtained after shuffling speed labels across trials, illustrating chance-level decoding structure.

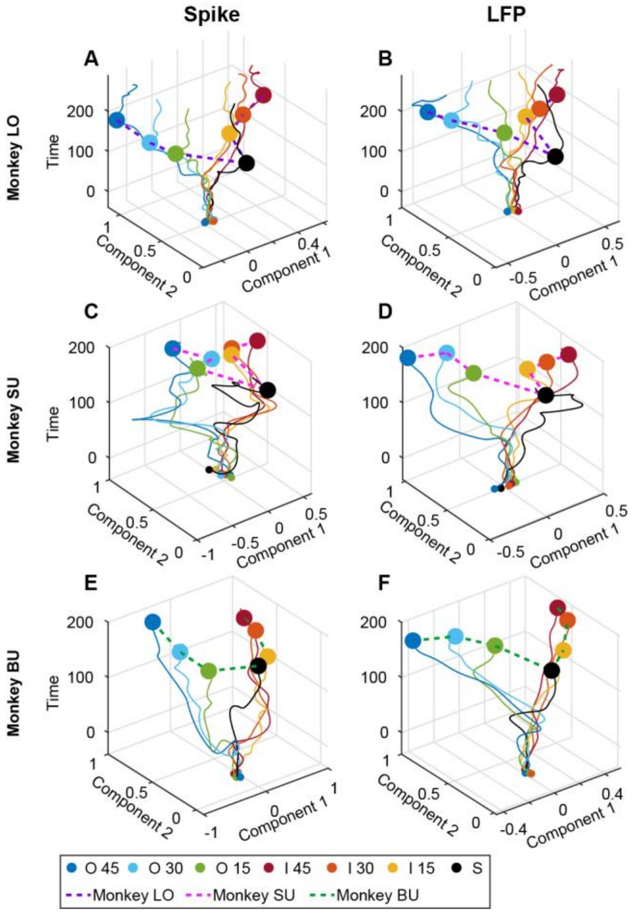


**Supplementary Figure 7.** Temporal evolution of SC population activity in reduced dimensions for the other three animals (LO: **A**, Spike; **B**, LFP; SU: **C**, Spike; **D**, LFP; BU: **E**, Spike; **F**, LFP). Trajectories depict population activity projected onto the first two LDA components, with time from target onset represented along the z-axis. Colors indicate speed condition. Large markers denote the 180ms time point. Same analysis as in Figure 6E & F.

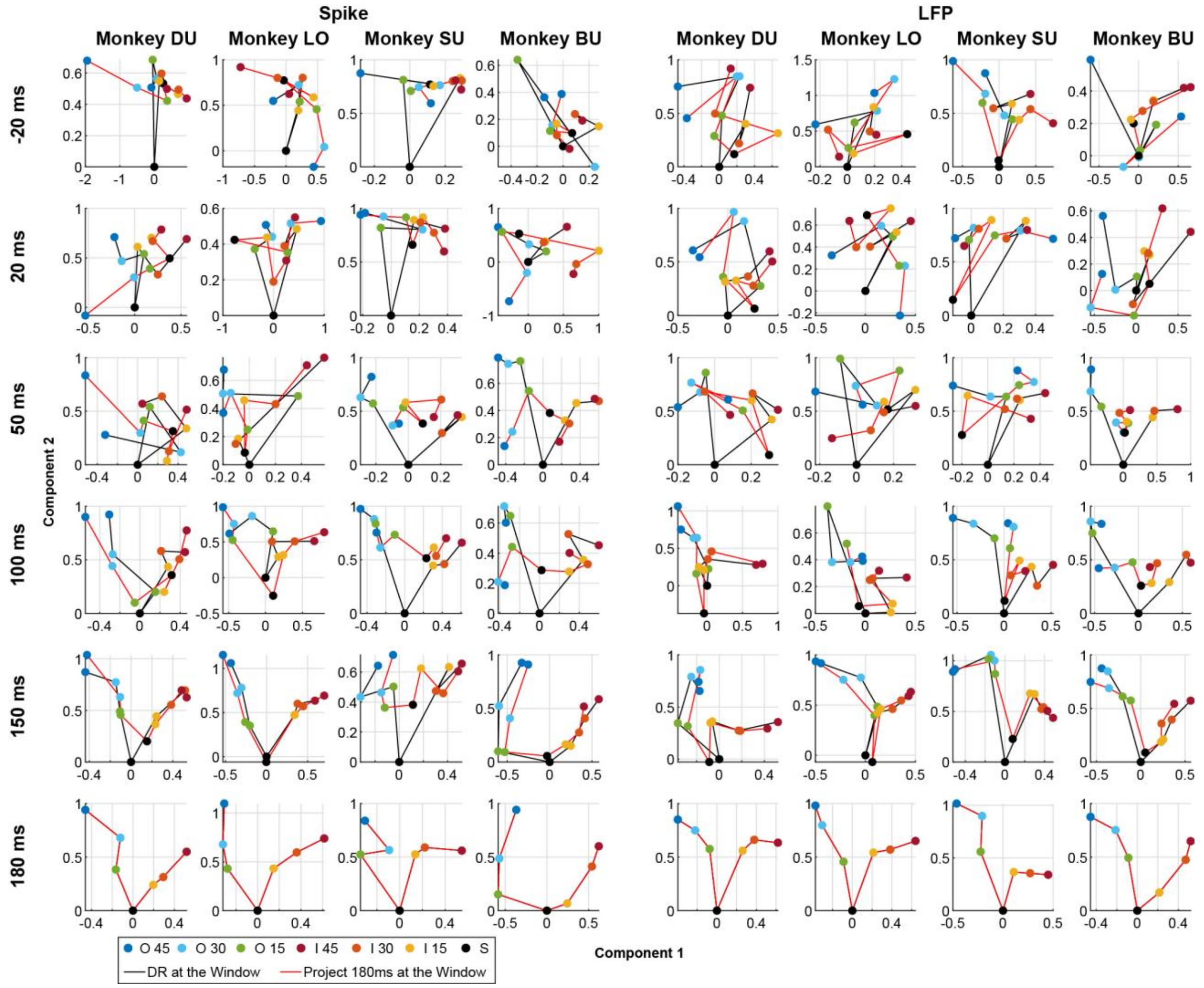


**Supplementary Figure 8.** Low dimensional population structure of target speed representation across time varying subspaces. Population activity projected into the first two dimensions defined by LDA. Columns show data from four monkeys (DU, LO, SU, BU), with spike activity on the left and LFP signals on the right. Rows indicate the time windows at which dimensionality reduction is performed relative to target onset (−20, 20, 50, 100, 150, and 180 ms). Points denote the cluster center for each speed condition (as in Figure 6C & D). Black traces show the population geometry obtained when dimensionality reduction is performed at the corresponding time window. Red traces show population activity at 180ms projected into the subspace defined at each time window. Across monkeys and signal types, population responses evolve from a weakly structured configuration before or shortly after target onset to a more organized low dimensional manifold at later time points. This structure progressively adopts a V shaped geometry, with increasing separation between speed conditions. The consistency between time specific trajectories and projections into the late subspace indicates that the geometric organization is not solely driven by time varying subspace rotations but reflects a stabilizing representation of target speed in population activity.

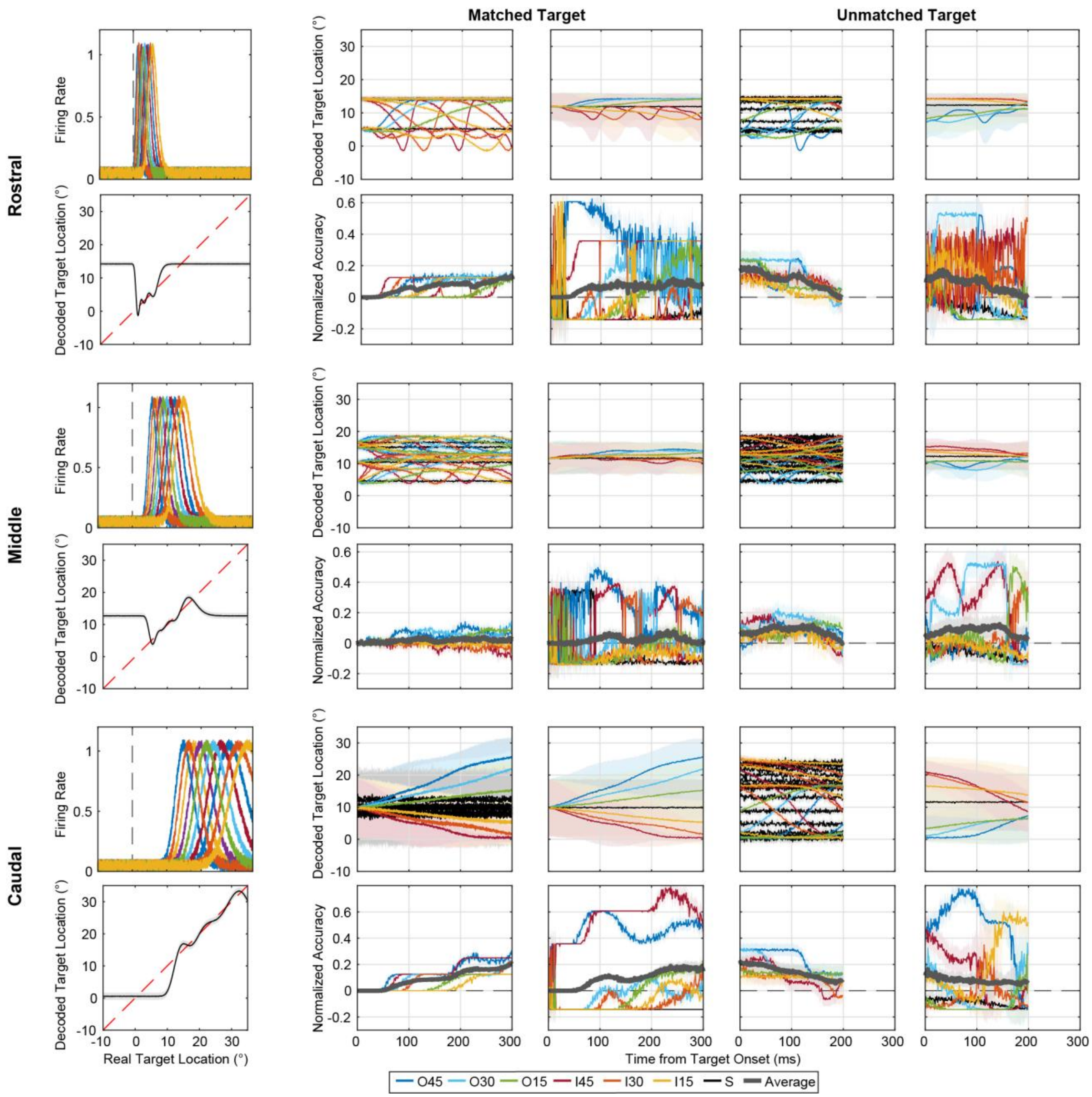


**Supplementary Figure 9**. Dependence of location based decoding simulations on the position of the sampled neuronal population within the SC map. Left column: Simulated neuronal receptive fields with noise (top, same as in **Figure 6 B**) and performance of the location decoder (bottom, same as in **Figure 6 C**) for populations centered at different locations along the SC map. Remaining columns: Simulated decoding results obtained using neuronal populations centered 1 mm (Rostral), 2 mm (Middle), and 3 mm (Caudal) from the rostral SC. Panels show decoded target location trajectories (same as in Figure 6 H-K) and decoding performance (same as in Figure 6 L-O) for matched and unmatched initial target configurations. For each recording location, decoded target trajectories are shown for individual target conditions and after averaging across initial target locations. Binary and multiclass decoding performance were. Colors denote target speed categories, and the gray trace indicates the average across conditions. Shaded regions indicate mean ± SD across simulated trials. Decoding accuracy is expressed as normalized accuracy (actual minus shuffled), with zero indicating chance performance.

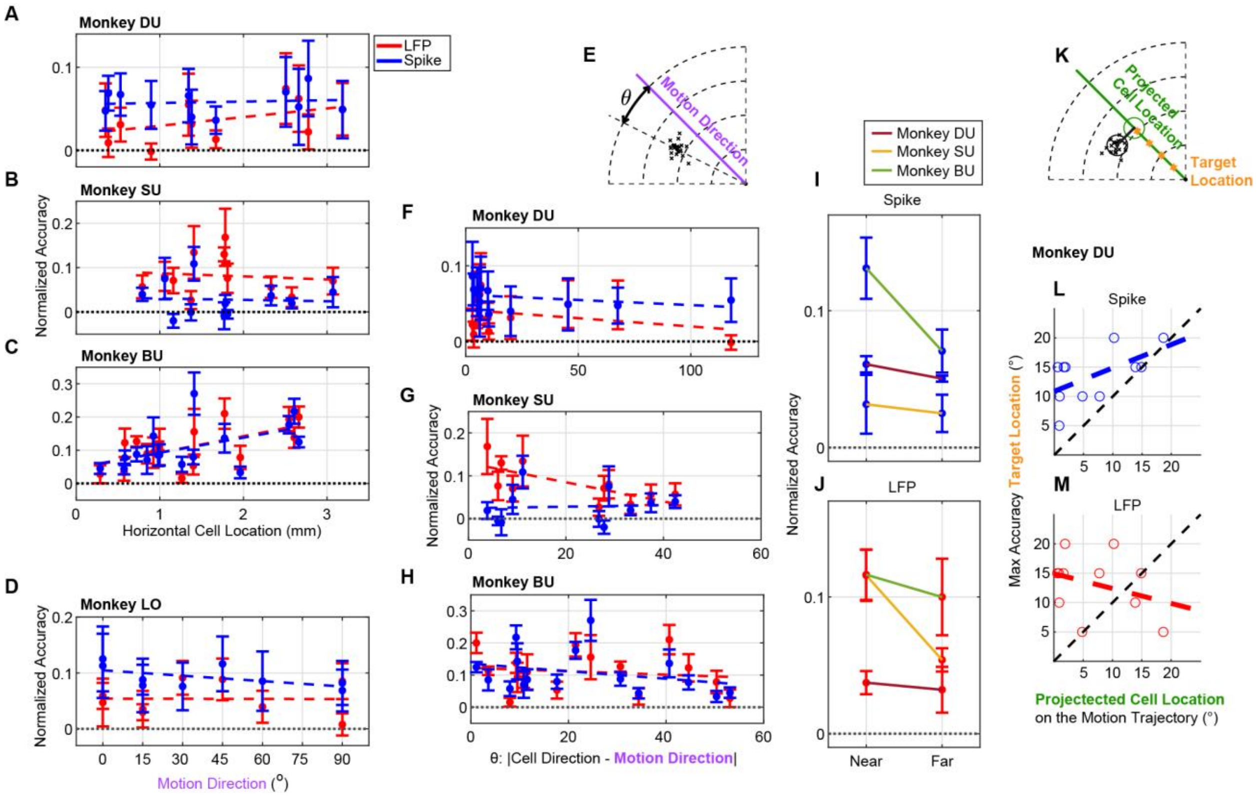


**Supplementary Figure 10.** Impact of cell location within SC on decoder performance. **(A-C)** Normalized decoding accuracy as a function of horizontal cell location for monkeys DU **(A)**, SU **(B)**, and BU **(C)**. The black dashed line indicates chance level (0). Each point represents a recording site. Blue color denotes spike-based decoding and red color denotes LFP-based decoding. Error bars indicate mean ± SD. Colored dashed lines show linear fits. **(D)** Normalized decoding accuracy as a function of target motion direction for monkey LO. Conventions are the same as in **(A-C)**. Note that data collected from the N-form array does not avail itself of repeating the analyses performed with laminar probes. **(E)** Schematic illustrating θ, defined as the angular difference between the preferred direction (center of mass across all cells recorded in a session) and the motion direction. **(F-H)** Normalized decoding accuracy plotted against |θ| (absolute angular difference between preferred and motion direction) for monkeys DU **(F)**, SU **(G)**, and BU **(H)**. Conventions are the same as in **(A-C)**. **(I** & **J)** Summary across monkeys (DU, SU, BU from **F-H**) comparing decoding accuracy for small ($|\theta| < 25°$) and large ($|\theta| \geqslant 25°$) differences in directions for Spike **(I)** and LFP **(J)**. **(K)** Schematic illustrating projected cell location along the motion trajectory relative to target location. **(L & M)** Relationship between the initial target location yielding maximum decoding accuracy and the projected cell location along the motion trajectory for spikes **(L)** and LFP **(M)** in monkey DU. Colored dashed lines indicate linear fits.
